# CRISPR-Cas-based Microhomology-Mediated End Joining for Precise Gene Replacement in Plant

**DOI:** 10.1101/2022.08.27.505510

**Authors:** Tien Van Vu, Gah-Hyun Lim, Seung Hee Choi, Ju Yeon Moon, Ngan Thi Nguyen, Swati Das, Mil Thi Tran, Yeon Woo Sung, Jihae Kim, Young Jong Song, Suk Weon Kim, Jae Cheol Jeong, Jae-Yean Kim

## Abstract

Precise gene or allele replacement is a desirable technology, but implementing it in plants remains challenging. CRISPR-Cas-based approaches, such as gene targeting (GT) and prime editing (PE), have opened up new possibilities for precise gene replacement in plants. However, their editing size and efficiency still need improvement. Recently, strategies using canonical nonhomologous end-joining (cNHEJ) and microhomology-mediated end joining (MMEJ) have been considered promising alternatives for precise gene replacement in yeast and mammals. However, these approaches have not been extensively explored and applied to plants. Here, we proposed and tested a CRISPR-Cas-based tool, termed PREMJ (<u>p</u>recision gene <u>re</u>placement via <u>m</u>icrohomology-mediated end joining), for precision gene replacement in plants. The PREMJ strategy employing 20-bp microhomology MMEJ donors (∼100 bp lengths) and a canonical nonhomologous end-joining (cNHEJ) inhibitor, NU7441, produced high targeted gene replacement efficiencies, up to 1.60 ± 0.14, 4.47 ± 1.98, and 8.98 ± 4.73 % in protoplasts of tomato, lettuce, and cabbage, respectively. Our data also revealed the critical impacts of the microhomology length and NU7441 concentration on PREMJ-based precision gene editing in plants. Although obtaining edited plants remains challenging due to inefficient protoplast regenerations and *Agrobacterium*-mediated delivery, PREMJ may significantly contribute to precision gene editing in plants competent to PREMJ complex delivery and plant regeneration.

**Key Message:** We designed and tested a method, termed PREMJ, for precise gene replacement using the CRISPR-Cas-mediated double-stranded break (DSB) formation and repairing the DSBs via microhomology-mediated end joining (MMEJ) with MMEJ donor template carrying desired base changes and microhomologies flanking the DSB ends. PREMJ showed feasibility in precise gene replacement in protoplasts of tomatoes, lettuce, and cabbage, albeit its efficacy in *Agrobacterium*-mediated plant transformation requires further optimization.

## Introduction

Precise modification of bases/genes of interest within their genomic context is highly desirable for gene functionalization and genetic engineering applications. However, efficient precision gene editing remains challenging in plants (Schindele et al. 2020). With the advent of the clustered regularly interspaced short palindromic repeats (CRISPR)/CRISPR-associated (Cas) technology (Jinek et al. 2012), plant precision gene editing has been revolutionized with homology-directed gene targeting (HGT) (Cermak et al. 2015; Wang et al. 2017; Wolter and Puchta 2019; Zhang et al. 2022; Vu et al. 2020) and prime editing (PE) (Anzalone et al. 2020; Lin et al. 2021; Vu et al. 2024b). With HGT, we can precisely edit genes from a single base to kilobase scales, while narrower-range precise gene editing could be achieved with the PE editing tools. However, HGT still requires improvement for practical applications to a wide range of *Arabidopsis* loci (Zhang et al. 2022) and crop plant loci due to low editing efficiency and inaccessibility of the genomic contexts of the loci (Wang et al. 2017; Vu et al. 2021a; Zhang et al. 2022). In the case of prime editors, while substantial improvements have been made to achieve efficient editing of short edits (Lin et al. 2021; Zong et al. 2022; Lu et al. 2020b; Vu et al. 2024b), precise insertion of DNA sequences with more than 50-bp lengths remains challenging (Vu et al. 2024a).

Double-stranded breaks (DSBs) formed by the activity of fully functional Cas9 proteins may be repaired majorly by canonical nonhomologous end-joining (cNHEJ) and homologous recombination (HR), and the former is the dominant pathway (Puchta 2005). MMEJ, previously thought to be an alternative pathway of the cNHEJ, has been shown to play a more important role in DSB repair (Tan et al. 2020; Ata et al. 2018; Sharma et al. 2015; Vu et al. 2021b). Recently, cNHEJ and microhomology-mediated end joining (MMEJ) mechanisms were repurposed for CRISPR/Cas9-based precise gene replacement at high efficiency and specificity in yeast and animals (Sakuma et al. 2016; Yao et al. 2017; Ata et al. 2018; Hayashi and Tanaka 2019; Li et al. 2019; Matsuzaki et al. 2024). In plants, Lu and co-workers engineered a cNHEJ-mediated CRISPR/Cas9-based precise gene insertion at high efficiency by employing a single cut at genomic sites in rice (Lu et al. 2020a). However, a system exploiting cNHEJ and MMEJ for precise gene replacement has not been developed in plants.

Here, we experimentally tested our previously proposed MMEJ-mediated precise gene editing (Vu et al. 2021b) and evaluated a novel MMEJ-based system for precise gene replacement, termed PREMJ, in plant protoplasts. Using PEG-mediated protoplast transfection and targeted deep sequencing methods, we successfully obtained highly precise editing efficiency in tomatoes, lettuce, and cabbage. The MMEJ-based precise gene replacement could be further improved by optimizing the microhomology lengths, donor template concentration, and suppression of cNHEJ by NU7441. However, PREMJ remained challenging with *in planta Agrobacterium*-mediated transformation methods, possibly due to the inefficiency in delivering sufficient amounts of MMEJ donors. Nevertheless, this tool may still offer an alternative way for precise gene editing in plants that facilitates efficient PREMJ tool delivery and regeneration and thereby help to advance crop breeding in the future.

## Materials and methods

### Donor DNA preparation for RNP works

Chemically modified donors (Supplemental Table 1; Supplemental Supplemental file 1) were prepared by annealing of synthesized oligos or by PCRs. The 5’ end modified cNJ.HPAT3-1 was prepared by PCRs using 5’ modified oligos (5’ phosphorylated, phosphorothioate bond addition to the phosphodiester linkage between the first and the second nucleotides) (Supplemental Table 1) synthesized by Bioneer (Korea) without a template. The MMEJ donors were prepared by PCRs using complementary oligos (Supplemental Table 1) synthesized by Bioneer (Daejeon, Korea) without templates. A high-fidelity DNA Taq polymerase was used for the PCR amplifications. The PCR products were cleaned using a BIOFACT PCR cleanup kit (BIOFACT, Korea). The donor concentrations were assessed by Nanodrop2000 spectrophotometer (Thermofisher, USA) and directly used for transfection stored at -20°C for further uses.

### Construction of plasmid for *Agrobacterium*-mediated transformation in tomato

For stable transformation and assessment of the cNHEJ and MMEJ-mediated precision gene replacement in tomatoes, we designed and cloned the gRNA expression cassettes (Supplemental file 1) using the Golden-gate cloning system as described previously (Vu et al. 2020). Two gRNA expression cassettes (gR1.HPAT3 and gR2.HPAT3 for *SlHPAT3*; gR1.HKT1;2 and gR2.HKT1;2 for *SlHKT1;2*) were used to generate two DSBs at each targeted site. The cNHEJ donors (cNJ.HPAT3-1 for SlHPAT3; cNJ.HKT1;2 for *SlHKT1;2*) and MMEJ donor (MJ.HPAT3-1 for SlHPAT3; MJ.HKT1;2 for *SlHKT1;2*) were designed to be flanked by two gRNAs (gDR1.HPAT3 and gDR2.HPAT3 for cNJ.HPAT3-1; gDR3.HPAT3-1 and gDR4.HPAT3 for MJ.HPAT3; gDR1.HKT1;2 and gDR2.HKT1;2 for cNJ.HKT1;2; gDR3.HKT1;2 and gDR4.HKT1;2 for MJ.HKT1;2) cutting sites (Supplemental file 1). The binary plasmids were constructed to test both the cNHEJ and MMEJ approaches using a conventional T-DNA and a geminiviral replicon system (Vu et al. 2020). The NptII selection marker expression cassette (pNOS-NptII-tOCS) is driven by the NOS promoter and terminated by the OCS terminator (Addgene # 51144). An intron-containing plant codon-optimized SpCas9 driven by a CaMV 35S promoter, and CaMV 35S terminator (p35S-pcoCas9I-t35S) was used (Supplemental file 1).

### Isolation of protoplasts

Tomato (Solanum lycopersicum cv. Micro-Tom), lettuce (Lactuca sativa L. cv. Cheongchima), and cabbage (Brassica oleracea) seeds were sterilized with 70% ethanol for 3 min, 1% hypochlorite solution for 15 min, and washed five times with distilled water. The sterilized seeds were inoculated in a medium containing 1/2 Murashige and Skoog salts, 0.4 mg/L thiamine HCl, 100 mg/L Myo-inositol, 30 g/L sucrose, and 8 g/L gelrite, pH 5.7. The seedlings were grown in a growth chamber under a 16 h light/8 h dark photoperiod (100–130 μmol/m2 s) at 25°C for tomato, and 20oC for lettuce, and 23oC for cabbage.

For protoplast isolation of tomato and cabbage, the cotyledons of 4-day-old tomato seedlings and the cotyledons of 7-day-old cabbage seedlings were immersed in cell and protoplast washing solution (CPW) containing 0.5% cellulase (Novozymes, Basgsvaerd, Denmark), 0.5% pectinase (Novozymes, Bagsværd, Denmark), 1% viscozyme (Novozymes, Bagsværd, Denmark), 3 mM MES (pH 5.8) and 9% mannitol. After 15 min of vacuum infiltration, the suspension was incubated for 2-4 hr on a rotary shaker at 50 rpm at 25°C. The suspension was filtered through an eight-layer gauze and centrifuged for 5 min at 100 *g*. Protoplasts were separated on a 21% sucrose density gradient and then collected at the interface of the W5 solution (2 mM MES pH 5.8, 154 mM NaCl, 125 mM CaCl_2_, 5 mM KCl). The harvested protoplasts were washed three times with W5 solution and then resuspended in MMG solution (4 mM MES pH 5.7, 0.4 M mannitol, 15 mM MgCl_2_). The concentration of protoplasts was determined using a hemocytometer.

For the lettuce protoplast isolation, the cotyledons of 7 d-old seedlings were digested with 10 mL of enzyme solution (1% [w/v] Viscozyme (Novozymes, Bagsværd, Denmark), 0.5% Celluclast (Novozymes, Bagsværd, Denmark), and 0.5% Pectinex (Novozymes, Bagsværd, Denmark), 3 mM MES (2-[N-Morpholino] ethanesulfonic acid), pH 5.7 and 9% mannitol in CPW salts with shaking at 40 rpm for 4–6 h at 25 °C in the dark. The protoplast mixture was then filtered through a 40 µm nylon cell strainer and collected by centrifugation at 800 rpm for 5 min in a 14 mL round tube. The collected protoplasts were re-suspended in W5 solution (2 mM MES [pH 5.7], 154 mM NaCl, 125 mM CaCl_2_, and 5 mM KCl) and further centrifuged at 800 rpm for 5 min. Finally, the protoplasts were re-suspended in W5 solution and counted under a microscope using a hemocytometer. Protoplasts were adjusted to a 1 × 106/mL density in MMG solution before transfection. The transfected protoplasts were cultured in protoplast culture medium (MS medium containing 0.4 mg/L thiamine HCl, 100 mg/L Myo-inositol, 30 g/L sucrose, 0.2 mg/L 2,4-dichlorophenoxyacetic acid [2,4-D], and 0.3 mg/L 6-benzylaminopurine [BAP], pH 5.7) in the dark f at 25℃ for 4 weeks.

### Protoplast transfections

SpCas9 protein was purchased from ToolGen, Inc. (Seoul, South Korea), and guide RNAs were synthesized by GeneArt Precision gRNA Synthesis Kit (Invitrogen, Massachusetts, USA) according to the manufacturer’s protocol. PEG-mediated RNP and donor transfections were performed in the previous study (Woo et al. 2015).

For RNP and donor DNA transfections with tomato and cabbage protoplasts, 2 × 10^5^ protoplasts were transfected with the purified SpCas9 protein (20 μg) premixed with sgRNAs (10 μg each) and donor templates in PBS buffer followed by incubating for 10 min at 25°C. The RNP complexes were mixed with protoplasts and then supplemented with an equal volume of 40% PEG transfection solution (40% PEG 4000, 0.2 M mannitol, and 0.1 M CaCl_2_). This suspension was mixed gently and then incubated at room temperature for 10 min. An equal volume of W5 solution was added for washing, followed by centrifugation at 100 *g* for 5 min. The supernatant was discarded, and the protoplasts were incubated with 1 ml of W5 solution in the dark at 25°C for 48 h. Afterward, the cells were collected for gDNA isolation and subsequent targeted deep sequencing analysis.

For lettuce protoplast transfection, SpCas9 protein and sgRNAs were premixed in 1× NEB buffer 3 for at least 10 minutes at room temperature, and 2×10^5^ protoplast cells were transfected with SpCas9 protein (20 μg) premixed with sgRNAs (10 μg each) and donor DNA. A mixture of 2 × 10^5^ protoplast cells was re-suspended in 200 μl MMG solution and then was slowly mixed with RNP complex, donor, and 350 μl of PEG solution (40% [w/v] PEG 4000, 0.2 M mannitol, and 0.1 M CaCl2). After incubation for 10 min, the transfected protoplast cells were gently re-suspended in 650 μL W5 solution. After additional incubation for 10 min, 650 μL W5 solution was added slowly again and was mixed well by inverting the tube. Protoplasts were pelleted by centrifugation at 556 rpm for 5 min and washed gently in 1 ml W5 solution. Protoplasts were pelleted by centrifugation at 556 rpm for 5 min and re-suspended gently in 1 ml WI solution (4 mM MES [pH 5.7], 0.5 M mannitol, and 20 mM KCl). Finally, the protoplasts were transferred into a 60 × 15 mm petri dish (Falcon), cultured under dark conditions at 25°C for 48 h, and then analyzed for genome editing efficiency.

### *Agrobacterium*-mediated tomato transformation

The *Agrobacterium*-mediated tomato transformation was conducted using a protocol published by Vu and coworkers (Vu et al. 2020) with or without 1 µM NU7441 treatment for 5 days post-washing. Ten days post-transformation, thirty cotyledon explants were collected per transformation plate to isolate genomic DNAs and subsequent miniseq analysis. Regenerated plants were selected in media containing 100 mg/L kanamycin and transferred to soil pots before analyzing for editing performance. Genomic DNAs were extracted from the plants and analyzed by PCR amplification of the targeted sequences and by Sanger sequencing.

### Targeted deep sequencing

Genomic DNAs were isolated from the protoplasts using the CTAB method. We used the miniseq sequencing service (MiniSeqTM System, Illumina, USA) to obtain targeted deep sequencing of the edited genomic sites. Miniseq samples were prepared in three PCRs according to the manufacturer’s guidelines, with genomic DNAs as the first PCR template. The first and second PCRs used primers listed in Supplemental Table 1, whereas the third PCRs were conducted with the manufacturer’s primers to assign a sample ID. A high-fidelity DNA Taq polymerase (Phusion, NEB, USA) was used for the PCRs. The miniseq raw data FASTQ files were analyzed using the Cas-Analyzer tool (Park et al. 2016). The indel examining window was set at 5 bases with a comparison range that covered both the read ends. A similar analysis was conducted for the targeted base changes of lettuce and cabbage genes.

### Statistical analysis

The editing data, statistical analysis, and plots were further processed by the MS Excel and GraphPad Prism 9.0 programs and are explained in detail in the legends of figures and tables wherever applicable. Statistical analysis was performed using a two-sided, uncorrected Fisher’s LSD test. A p-value of 0.05 or lower is considered statistically significant.

## Results and discussion

### System design and validation for precise gene replacement in plants

In a PREMJ system design, a predefined genomic site with a protospacer adjacent motif (PAM) is determined first, thus enabling the prediction of the DSB formation site (Figure 1A and 1B). The sequence between the two predicted cut sites is employed as a donor template so that its DNA sequence is essentially modified, allowing it to carry targeted base changes to avoid recurrent cuts after gene replacement (Figure 1A, 1B, and Supplemental Figure 1). In the case of MMEJ-mediated editing, the predicted flanking DSB ends would then be used to choose microhomologies (MH1, and MH2, Supplemental Figure 1B) at different lengths. Still, they should be optimally at least 8 nt (up to 20 nt was reported) in size (xxxxxxxx and yyyyyyyy in Figure 1B), as suggested (Nakade et al. 2014; Sakuma et al. 2016; Ata et al. 2018).

**Figure 1.**
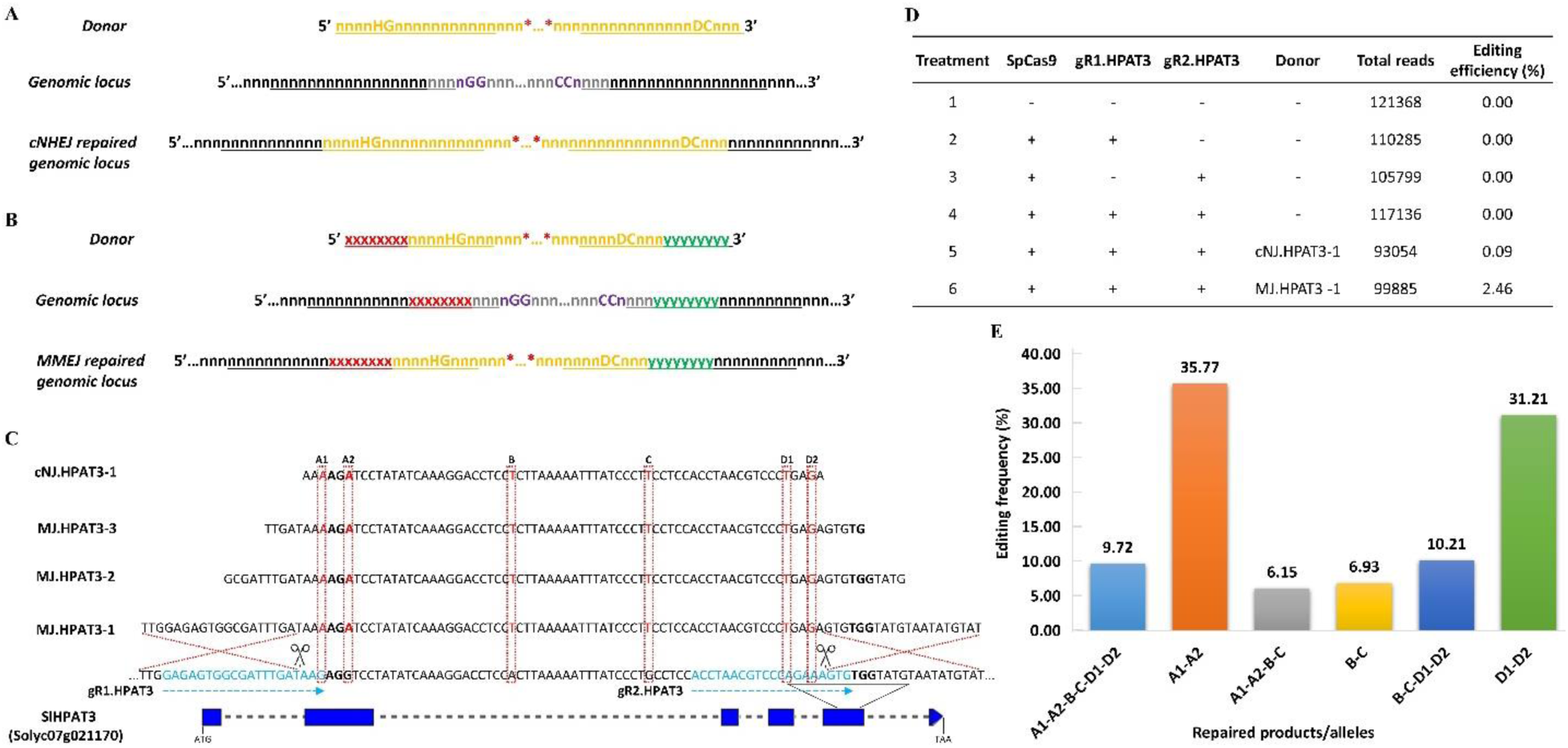
HR-independent precision editing approaches using CRISPR/SpCas9. (**A**) Model of sequence design for cNHEJ-based precise gene replacement. (**B**) Model of sequence design for MMEJ-bases precise gene replacement. The cut sites are arranged to avoid recurrent cutting. Active PAM sites are shown in purple. Guide RNA sequences or traces of them in repaired products are underlined. The red xxxxxxxx represents MH1, whereas the green yyyyyyyy represents MH2. The orange sequences denote donors, and the gray indicates the genomic sequence being replaced. The red asterisks represent sequence modifications of interest. The letters n, x, and y represent any nucleotide of A, T, G, or C, except that the sequence-making gRNA spacers should not have more than 4 T in a row to avoid early transcription termination. To prevent the recurrent cut, the PAM sequence corresponding to the replaced genomic sequence should be changed. Usually, nGG/CCn is changed to nHG/CDn where H=A or C or T; D=A or G or T. One or some bases within the core spacer sequence (i.e., ∼12 base pairs upstream/downstream of nGG/CCn) could also be changed. Such changes should not lead to changes in protein sequences of the targeted gene. (**C**) Schematic diagrams of cNHEJ- and MMEJ-mediated gene editing of tomato targeted locus *SlHPAT3* (with their GenBank accession number), targeted sequences, gRNA binding sites, and donor sense sequences (cNJ.HPAT3-1, MJ.HPAT3-1, MJ.HPAT3-2, and MJ.HPAT3-3). The targeted genes are shown from the start codon (ATG) to the stop codon (TAA), exons are drawn as blue boxes, and the discontinuous lines in each gene represent intron parts of the gene; the gRNA binding sequences at the genomic locus are highlighted in light blue color and arrows with PAMs (bold nucleotides). The expected cutting sites (3-nt upstream of a PAM) are indicated with the black scissors; the intended base changes are painted in the red font that is denoted by the discontinuous red boxes and named A1, A2, B, C, D1, and D2 on the top of the boxes. The cNHEJ donor dsDNA is synthesized with 5’-phosphorylated-ends, and the phosphodiester bonds between the first and the second nucleotides of 5’ ends were chemically modified with phosphorothioate linkages. The two red, discontinuous crossing lines indicate the MMEJ-mediated replacement sites of the MJ.HPAT3-1 donor. For the donors with shorter microhomology, the recombination lengths are reduced accordingly. The diagrams are not drawn to their scales. (**D**) cNHEJ and MMEJ-mediated editing efficiency at the SlHPAT3 locus. The editing efficiency is calculated by dividing the total reads containing the intended bases by the total reads without considering the indel mutation efficiency of the gRNAs. (**E**) Editing frequency of respective repaired products/alleles revealed from miniseq data. The edited base-changed sets/alleles revealed from miniseq data are classified according to the presence of the base changes in the edited reads. A total of 11 base-changed sets/alleles are recorded and plotted with their relative frequency among all the edited reads.

We selected a tomato homolog of *Arabidopsis* hydroxyproline O-arabinosyltransferase 3 (*SlHPAT3*, accession no. Solyc07g021170) for our proof-of-concept experiments (Brooks et al. 2014). We then employed PEG-mediated tomato protoplast transfection to deliver the SpCas9 proteins, two sgRNAs, namely gR1.HPAT3 and gR2.HPAT3 (Figure 1C) for cutting the genomic sites, and donors for replacing six base pairs of SlHPAT3 (Figure 1C; Supplemental Table 1 and Supplemental file 1). With the treatments of the SpCas9 protein and either of the gR1.HPAT3 or gR2.HPAT3, or both of them, without a donor, no precise edited products were found (Figure 1D). When either the NHEJ or the MMEJ donor was employed with both the gRNAs, precise replacement occurred, as shown by Sanger sequencing (Supplemental Figure 2 and Supplemental Table 2). Further analysis of the targeted deep-sequencing data revealed editing efficiency as high as 2.46 % (Figure 1D and Supplemental Table 3) in the case of the MJ.HPAT3-1 donor. Adding the cNJ.HPAT3-1 donor also resulted in the precise replacement of some bases pre-introduced in the donor sequence at a much lower frequency of 0.09 %. Significantly, the edited products of the cNHEJ donor did not include base changes located at its two ends (Supplemental Tables 2 and 3), indicating that either the donor terminal ends might be damaged before use for editing or that an unknown mechanism, possibly oligonucleotide-directed mutagenesis (ODM), might involve in the integration of the middle SNPs into the genomic sequence.

Further analysis of the edited products revealed multiple repaired products by the MJ.HPAT3-1 donor with various frequencies (Figure 1E). The precisely edited allele containing the intended base modification accounted for 9.72 % of total edited reads (Supplemental Table 3). However, when we observed all the edited products containing the desirable B and C base changes and excluded the others containing one-sided base changes (A1 and A2 or D1 and D2) that were designed to prohibit the recurrent cleavages of the SpCas9, the proportion of these alleles was 33.02% of the total edited alleles (Figure 1E; Supplemental Table 3). The diversity of the repaired products indicates that not only was MMEJ involved in repairing the DNA damage, but it also participated in other repair mechanisms like HR for one-sided alleles and ODM with only the B and C base changes. The initial data showed that both cNHEJ and MMEJ-mediated precision gene editing are feasible in tomato protoplasts, with the latter performing far better than the former.

### Higher donor amounts enhance PREMJ efficiency

The copy number of donor DNA templates available at the targeting sites during the cleavage of CRISPR/Cas complexes was shown to be essential for enhancing HDR efficiency (Cermak et al. 2015; Vu et al. 2020; Wang et al. 2017). We sought to investigate the implications of different donor doses on the frequency of PREMJ. The PREMJ frequencies with all base changes were increased with higher amounts of donor DNA, from 10.42 % (50 pmol) to 19.84 % (300 pmol), and the one-sided repair frequencies were reduced accordingly (Figure 2A and Supplemental Figure 5; Supplemental Table 4). In the case of the cNJ.HPAT3-1 donor, the editing efficiency did not vary much among the donor doses and was much lower than that of the MJ.HPAT3-1 (Figure 2A). Again, most of its products contained only B and C changes (Supplemental Figure 5). These data indicate that the simultaneity of the MMEJ donor presence and the cleavages of the genomic sites was important for precise gene replacement since higher donor doses increased the chance of diffusing them to the being-cleaved targeted sites. The higher abundance of the donor templates might also ensure sufficient amounts of undamaged donor DNA templates for editing.

**Figure 2.**
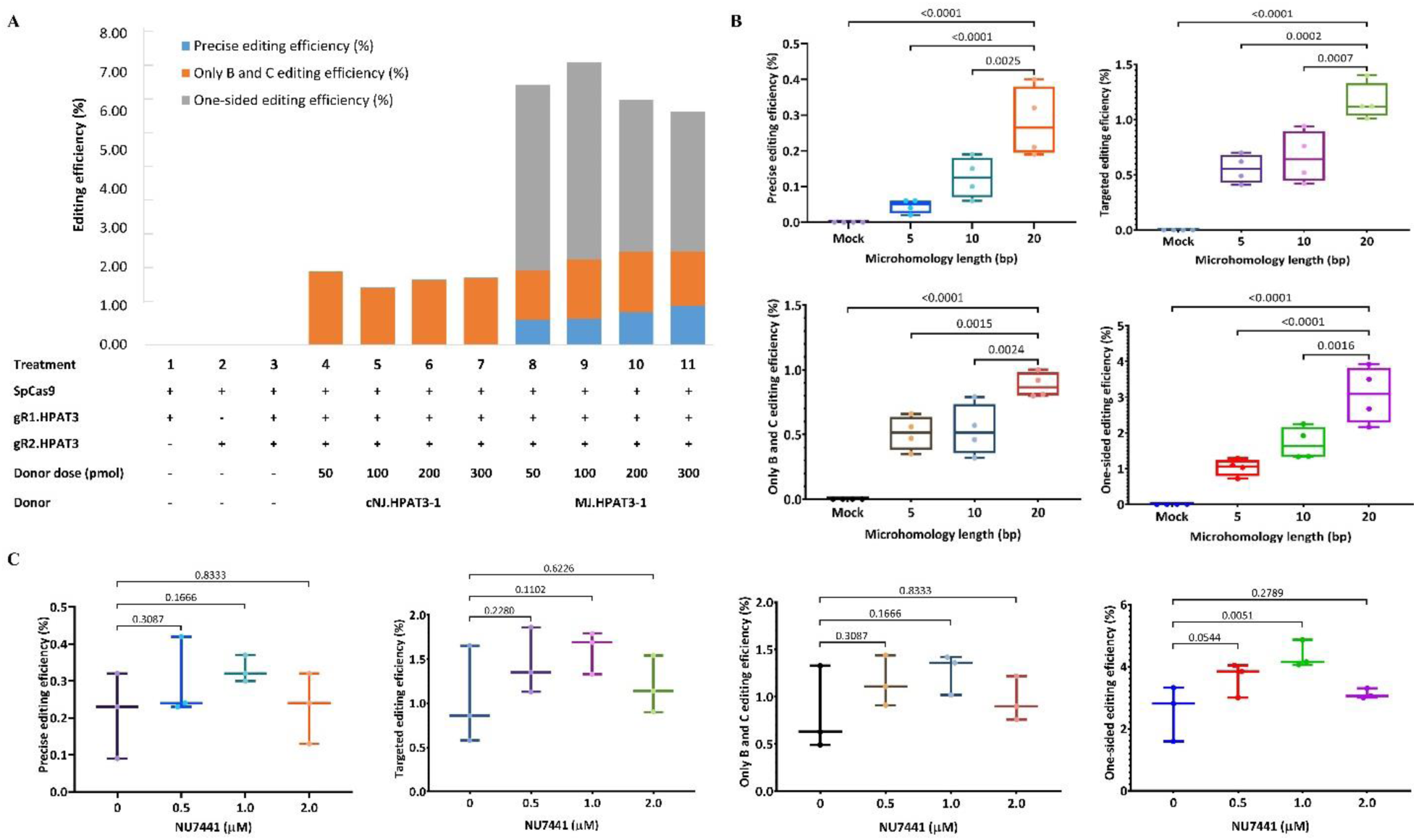
Optimization of MMEJ-mediated precision gene editing in tomato. **(A)** Editing efficiency revealed from targeted deep sequencing data of the various donor doses. **(B)** Editing efficiencies revealed from the treatments of donors with different microhomology lengths. Targeted deep sequencing data were collected in four replicates with no donor (-donor, Mock) control. **(C)** Editing efficiency revealed from targeted deep sequencing data of the treatments at different NU7441 concentrations. Targeted deep sequencing data were collected in three biological replicates. The data in (**B**) and (**C**) were analyzed by the Cas-Analyzer tool, statistically analyzed (multiple comparisons used a two-sided, uncorrected Fisher’s LSD test), and plotted by GraphPad Prism 9. The p values are shown on the top of each compared pair. All the data points are shown on the plots. Targeted editing efficiency is the sum of the precise editing efficiency and the only B and C editing efficiency. Precise editing efficiency was calculated by dividing the total reads containing all the base changes by the total reads. The only B and C editing efficiency was calculated by dividing the total reads containing only B and C base changes by the total reads. One-sided editing efficiency was calculated by dividing total reads containing only either the left-sided bases (A1 and A2) or the right-sided base pairs (D1 and D2) by the total reads.

### The length of microhomology affects PREMJ performance

The length of microhomology arms was an important parameter for activating the MMEJ pathway upon DSB formation and end resection (Villarreal et al. 2012). Effective microhomology lengths (8-20 bases) increased MMEJ-mediated gene insertion in mammalian cells (Nakade et al. 2014; Sakuma et al. 2016; Ata et al. 2018). We, therefore, tested the impact of three different microhomology lengths (20, 10, and 5 bp) on PREMJ efficiency. When the microhomology was shorter than 20 bp, the total editing efficiency was significantly reduced from 4.22 ± 0.47 % (MJ.HPAT3-1) to 2.33 ± 0.31 % (MJ.HPAT3-2) and 2.65 ± 0.58 % (MJ.HPAT3-3) (Supplemental Table 6). More importantly, the all-base-changed precise editing efficiency was significantly higher for 20-bp microhomology (0.28 ± 0.05 %) compared to that of the 10-bp (0.12 ± 0.03 %) and 5-bp (0.04 ± 0.01 %) microhomology lengths (Figure 2B). Furthermore, when we consider the reads, including the B and C base changes, the targeted editing efficiency was 1.16 ± 0.09, 0.66 ± 0.12, and 0.55 ± 0.06 % for the 20-bp; 10-bp, and 5-bp microhomologies, respectively (Figure 2B; Supplemental Table 6). Interestingly, the activities of the gRNAs were not significantly different among all the treatments (Supplemental Table 6), indicating that the intermolecular MMEJ-mediated editing efficiency obtained from the experiments depended on microhomology length, as shown in the cases of intramolecular MMEJ (Ata et al. 2018; Bae et al. 2014; Nakade et al. 2014). These data suggest that the 20-bp microhomology was the best among the tested lengths, confirming the highest performance of the 20-bp microhomology (Sakuma et al. 2016) for MMEJ-mediated DSB repair in plants.

### NU7441 improves PREMJ

Since the PREMJ efficiency was still low to be practically applicable for crop improvement, significant improvement of the editing system is needed. NU7441, a small chemical that was shown to inhibit DNA-dependent protein kinase (DNA-PKcs), an important cNHEJ component, enhancing HR-mediated repair of DSBs (Zhao et al. 2006; Robert et al. 2015; Vu et al. 2021a), also significantly elevated the MMEJ-mediated DSB repair products in mammalian cells (Dutta et al. 2017). We next tested if NU7441 can facilitate MMEJ repair using various concentrations of the chemical and the MJ.HPAT3-1donors. When the NU7441 concentration was increased from 0 to 1 µM, the editing efficiency was significantly elevated (Table 1 and Figure 2C). The read numbers of the precise products containing all the intended base changes, or only B and C base changes, were enhanced with the targeted editing efficiency reached up to 1.60 ± 0.14 % at 1 µM NU7441 compared to 1.03 ± 0.32 % without the addition of NU7441 (Table 1). The total editing efficiency was significantly higher with 1 µM NU7441 treatment, reaching up to 5.96 ± 0.34 %, a 1.65-fold increase compared to mock control and majorly contributed by one-side editing (Table 1), indicating that NU7441 treatment effectively favored the choice of MMEJ pathway for DSB repair. When NU7441 was added up to 2 µM, the repaired outcomes were reversely correlated with the NU7441 concentrations (Figure 2C). This result suggests that over-suppression of the cNHEJ components might lead to cellular toxicity, as shown in mammalian cells (Robert et al. 2015), and thus negatively impact the DSB repairs (Table 1 and Figure 2C).

**Table 1.**
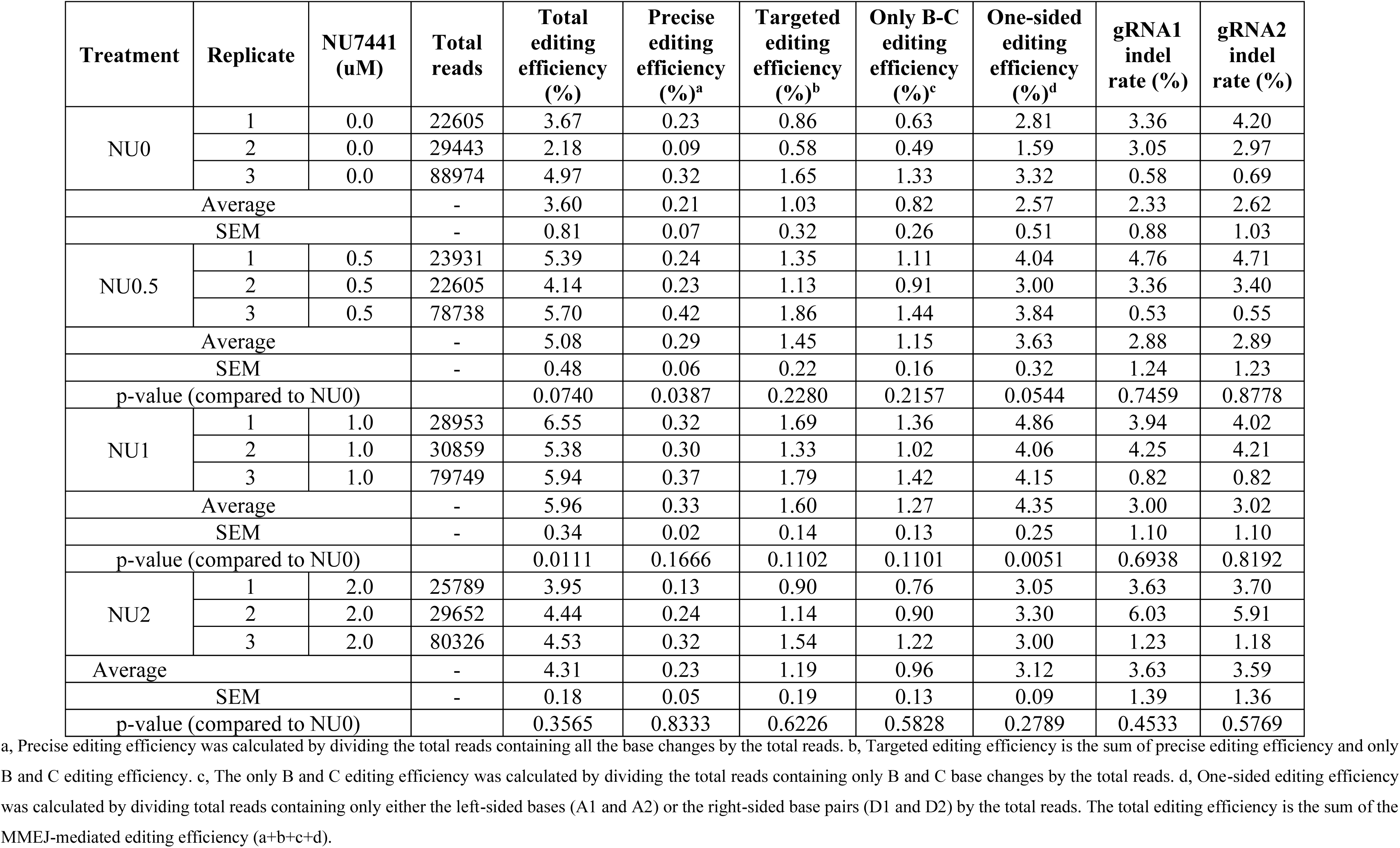
The detailed targeted deep sequencing data revealed from the various NU7441 concentrations.

Taken together, MMEJ-mediated precision editing is feasible in tomatoes and might be improved by using sufficient microhomology and NU7441 treatment.

### Efficient PREMJ-based gene editing in lettuce and cabbage protoplasts

To extend PREMJ-mediated precision gene editing to other plant species, we conducted PREMJ-mediated precise gene replacement in lettuce and cabbage protoplasts using the ribonucleoprotein (RNP) transfection method. The *THERMO-TOLERANCE 1* (*TT1*) (Li et al. 2015), *ORANGE* (*Or*) (Yuan et al. 2015; Yazdani et al. 2019), and *ACETOLACTATE SYNTHASE 1* (*ALS1*) (Mazur et al. 1987) genes were selected as targets for both lettuce and cabbage (Supplemental Figure 6). All the edited alleles of *TT1*, *Or*, and *ALS1* contained a single amino acid substitution (Supplemental Figure 6) for obtaining the expected traits (heat tolerance, carotenoid overaccumulation, and herbicide tolerance, respectively) (Li et al. 2015; Yuan et al. 2015; Yazdani et al. 2019; Mazur et al. 1987). Sequence alignment using tomato protein sequences was conducted using NCBI Blastp and putative *TT1*, *Or*, and *ALS1* genes were subsequently identified in lettuce (Supplemental Tables 7) and cabbage (Supplemental Tables 8). We designed and employed two gRNAs for cutting genomic loci and MMEJ donors containing 20-base microhomologies at two ends (Figure 3A and 3B). In lettuce, the highest precise PREMJ efficiency for all the intended base changes was 1.81 ± 0.75 % for the *LsALS1* locus, and the lowest efficiency was zero for the *LsTT1* locus. The *LsOr* locus showed only 0.13 ± 0.10 % for exchanging all the intended bases (Figure 3C; Supplemental Table 9). As mentioned earlier, the simultaneous activity of the two gRNAs is important for PREMJ-mediated DSB repair. The data here suggest more similar cleavages between the two gRNAs of *LsOr* and *LsALS1* compared to that of the *LsTT1* (Supplemental Table 9), which may partially explain the inefficiency of PREMJ-mediated repair at that locus. Considering all the repaired products that contain the targeted base changes for the desirable a.a., the efficiency reached 4.47 ± 1.98 %, 0.78 ± 0.30 %, and 0.42 ± 0.28 % for *LsALS1*, *LsTT1*, and *LsOr* locus, respectively (Fig 3C; Supplemental Table 9).

**Figure 3.**
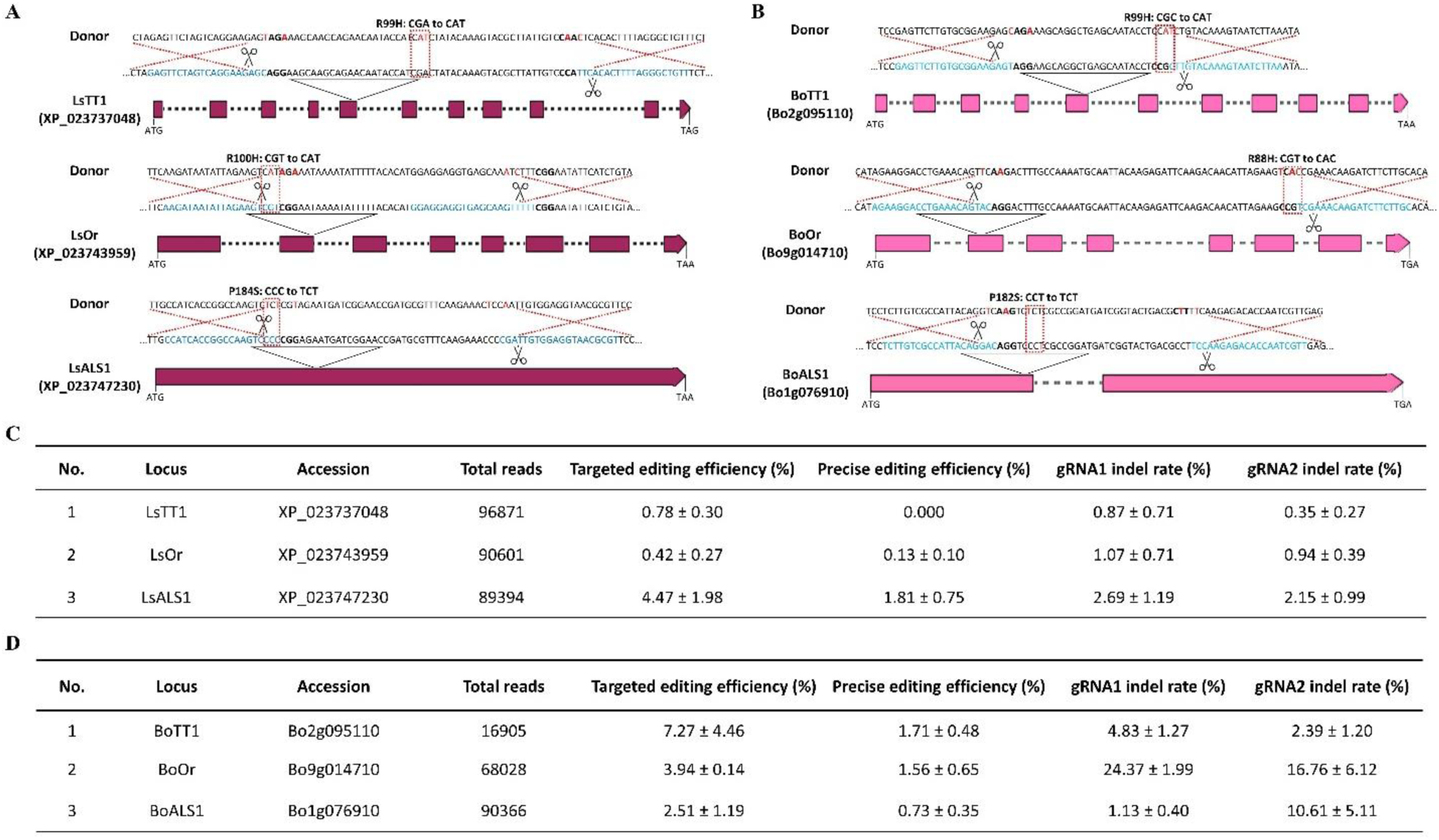
MMEJ-mediated precision gene editing in lettuce and cabbage. **(A-B**) Schematic diagrams of targeted loci (*TT1*, *Or*, and *ALS1* with their GenBank accession number), targeted sequences, gRNA binding sites, and MMEJ donors in lettuce (**A**) and cabbage (**B**). The targeted genes are shown from the start codon (ATG) to stop codons. Exons are drawn as colored boxes (purple boxes with lettuce’s loci and pink boxes for cabbage), and the discontinuous line in each gene represents intron parts of the gene; the gRNA binding sequences are highlighted in light blue color with PAMs (bold nucleotides). The expected cutting sites (3-nt upstream of a PAM) are indicated with the black scissors; the intended base changes are painted in red font; the targeted amino acid changes and corresponding triplexes are denoted by the discontinuous red boxes, and the texts placed on their top. The diagrams are not drawn to their scales. **(C**) MMEJ-mediated editing efficiency in lettuce at the *TT1*, *Or*, and *ALS1* loci. (**D**) MMEJ-mediated editing efficiency in cabbage at the *TT1*, *Or*, and *ALS1* loci. Targeted deep sequencing data were collected in three replicates and were analyzed by the Cas-Analyzer tool.

In cabbage protoplasts, the PREMJ worked more efficiently than for the tomato and lettuce protoplasts. The highest precise editing efficiency (7.27 ± 4.46 %) was obtained at the *BoTT1* locus and lower at the *BoOr* gene (1.68 ± 0.24 %). The *ALS1* locus for cabbage resulted in the least precise editing efficiency, at only 0.73 ± 0.43 % (Figure 3D; Supplemental Table 10). Moreover, the targeted editing efficiency at the *BoTT1* locus was the highest among tested loci in tomato, lettuce, and cabbage, reaching up to 8.98 ± 4.73 % (Figure 3D). In summary, the data from lettuce and cabbage indicate that PREMJ-mediated precision editing could be successfully extended to other plant species.

### *Agrobacterium*-mediated delivery of PREMJ in tomatoes

The plant regeneration system for tomato protoplast was shown to be of very low efficiency and time-consuming. To overcome this challenge, we attempted to deliver the PREMJ tools into tomato cotyledon explants by the *Agrobacterium*-mediated method (Vu et al. 2020). *SlHPAT3* and *SlHKT1;2* loci were selected as the targets for allele replacement (Supplemental Figure 7a,b and Supplemental file 1). Unexpectedly, at the callus stage, the PREMJ efficiency was extremely low at both the loci (Supplemental Table 11) and the editing reads were mostly with one-side editing (Supplemental Figure 8a) or one-sided insertion of the MMEJ donor (Supplemental Figure 8b). However, subsequent screening of transformants revealed plants carried precisely edited alleles, albeit the editing frequency was relatively low (up to 4% for the cNHEJ event cNJ1 and 3% for the PREMJ event #MJ5) (Supplemental Figure 9). The low efficiency of PREMJ might be explained by the requirement of simultaneous cleavages of four sites located in the donors and genomic sequences. In addition, the donor template amounts might also not be sufficient to be distributed to the targeted sites immediately after DSB formations. Those hurdles are not subjected to the protoplast transfection method.

## Conclusions

We have developed and validated the PREMJ method in plant protoplasts. The PREMJ approach has no limitation concerning the position and the number of base changes within the donor sequence. Such gene replacement is helpful in short sequence replacement for engineering regulatory elements, *in vivo* protein tagging, and short a.a. modification of a protein. Multiple-site editing is also possible with the RNP-protoplast transfection, but further testing is required. However, *in planta* transformation and regeneration efficiency using PREMJ tools should be sufficiently improved for practical applications in plant breeding.

## Data availability

All data generated or analyzed during this study are included in this published article and its supplementary information files or it will be provided upon a reasonable request.

## Acknowledgments

Not applicable.

## Author information

### Authors and Affiliations

Division of Applied Life Science (BK21 FOUR Program), Plant Molecular Biology and Biotechnology Research Center, Gyeongsang National University, Jinju 660-701, Republic of Korea

Tien Van Vu, Ngan Thi Nguyen, Swati Das, Mil Thi Tran, Yeon Woo Sung, Jihae Kim, Young Jong Song, Jae-Yean Kim

Department of Biological Sciences, Pusan National University, Busan 46241, Republic of Korea Gah-Hyun Lim

Biological Resource Center, Korea Research Institute of Bioscience and Biotechnology (KRIBB), Jeongeup, 56212, Republic of Korea

Gah-Hyun Lim, Seung Hee Choi, Ju Yeon Moon, Suk Weon Kim, Mil Thi Tran, Jae Cheol Jeong Nulla Bio Inc 501 Jinju-daero, Jinju 52828, Republic of Korea

Jae-Yean Kim

### Contributions

Conceptualization, T.V.V and J.Y.K.; Methodology, T.V. V and J.Y.K.; Conducted experiments, T.V.V., G.H.L, S.H.C, J.Y.M., S.W.K., J.C.J, N.T.N, S.D., M.T.T., Y.W.S, J.K, Y.J.S; Data analysis, T.V.V. and J.Y.K.; Writing – Original Draft, T.V.V.; Writing – Review & Editing, T.V.V. and J.Y.K.; Funding Acquisition, T.V.V., and J.Y.K.; and Supervision, T.V.V and J.Y.K.

## Ethics declarations

### Consent for publication

Not applicable.

### Ethical approval

Not applicable.

## Supplemental information

## SUPPLEMENTAL FIGURES

**Supplemental Figure 1.**
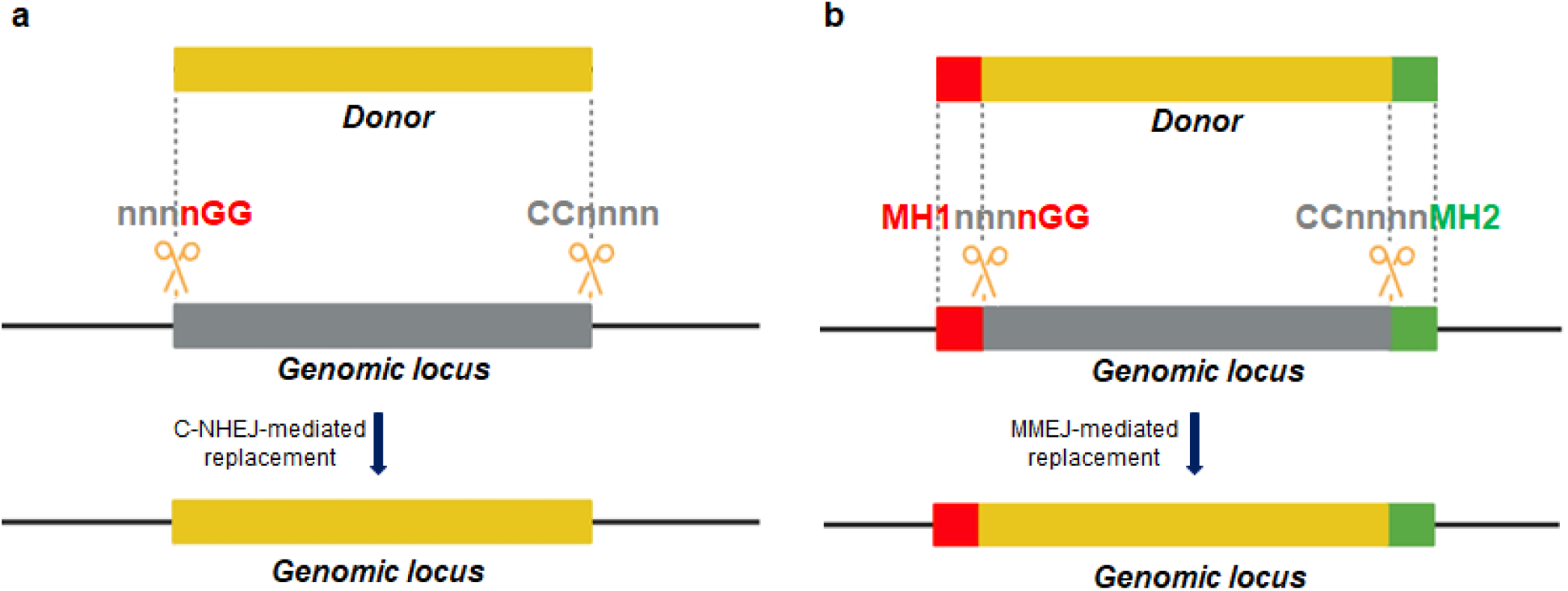
HR-independent strategies for precision gene/allele replacement using CRISPR/SpCas9. (**a**) cNHEJ-mediated DNA replacement model. (**b**) MMEJ-mediated DNA replacement model. MH1 and MH2 are predefined from genomic loci and are subsequently introduced into the two ends of the donor sequence. CRISPR/Cas complexes are designed with canonical PAM binding sites (nGG/CCn) to produce DSBs at genomic loci with MH1 and MH2 flanking ends. MMEJ-mediated repair using the donor sequence generates genomic loci with precise modifications. The cut sites are arranged to avoid recurrent cutting. The orange sequences denote donors, and the gray indicates the genomic sequence being replaced. The letters n represent any nucleotide of A, T, G, or C except that the sequence-making gRNA spacers should not have more than 4 T in a row to avoid early transcription termination.

**Supplemental Figure 2.**
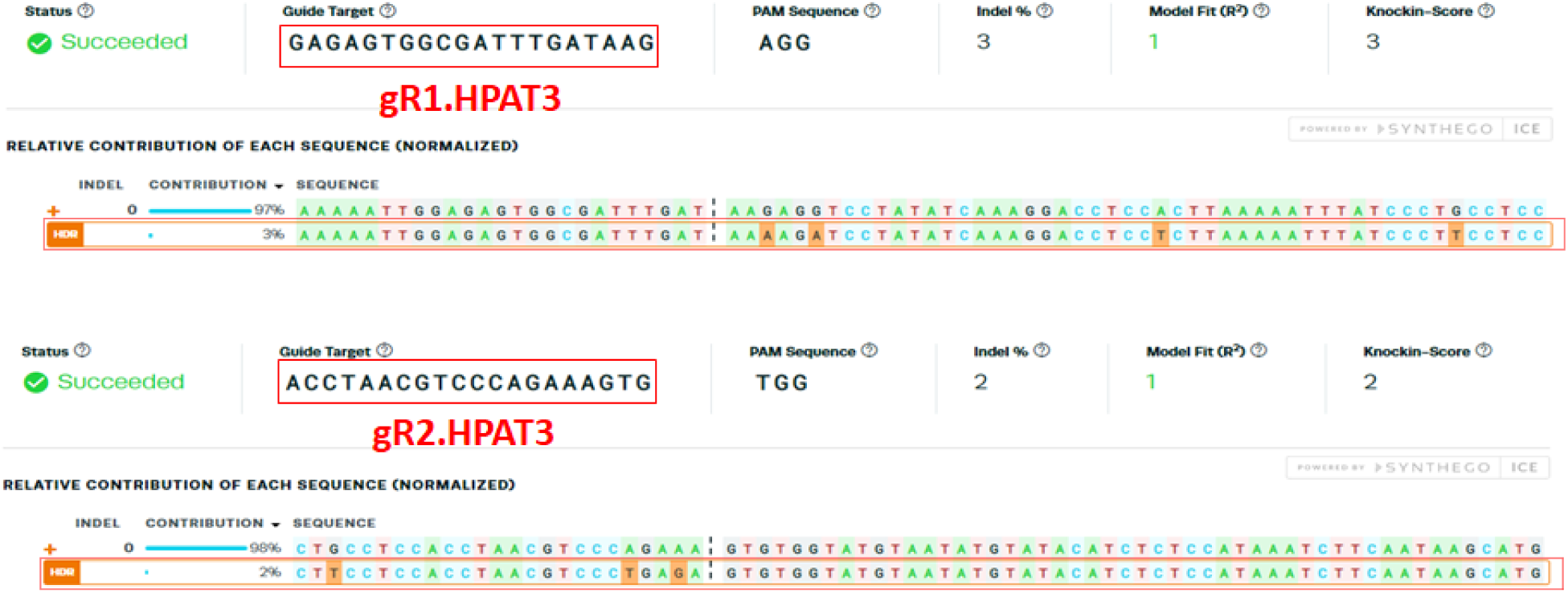
Sanger sequencing data revealed precise replacement of DNA by MMEJ donor. PCR amplification of the targeted site was conducted using primers flanking the site and PCR products were resolved on gel and purified for Sanger sequencing. Sanger sequencing data were decomposed using the ICE Synthego web application using gR1.HPAT3 (upper panel) and gR2.HPAT3 (lower panel) for locating the cutting site. The edited allele and its percentage among the total sequence were revealed and highlighted in the discontinuous red boxes. The indented base changes of the left and right sides of the targeted sequence are highlighted in the orange background of the up and down panels, respectively.

**Supplemental Figure 3.**
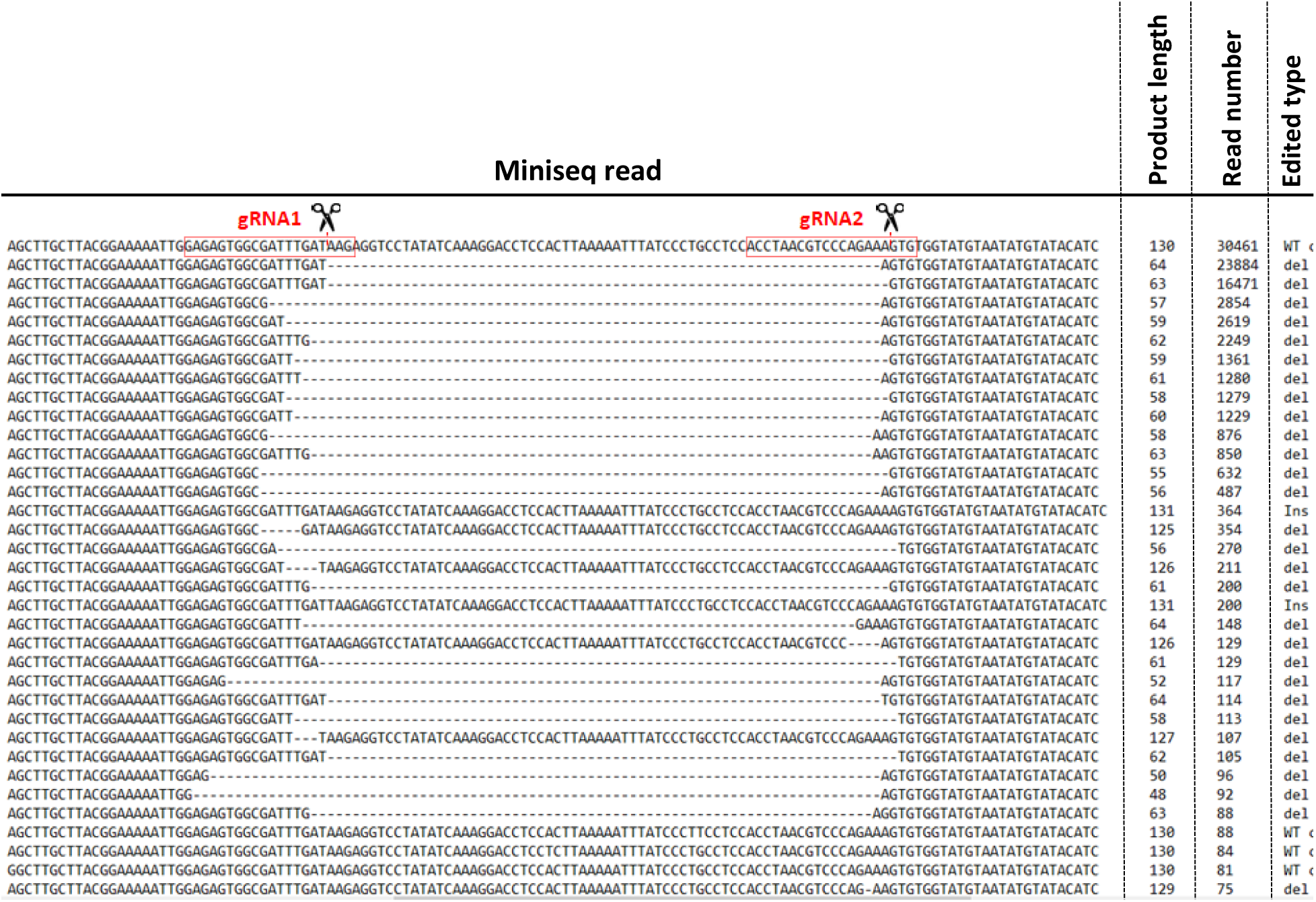
Indel alleles revealed from transfection of SpCas9, gR1.HPAT1 and gR2.HPAT3 with cNJ.HPAT3-1 donor. The gRNA binding and cutting sites are shown on the top panel with the scissor icons on the WT sequence.

**Supplemental Figure 4.**
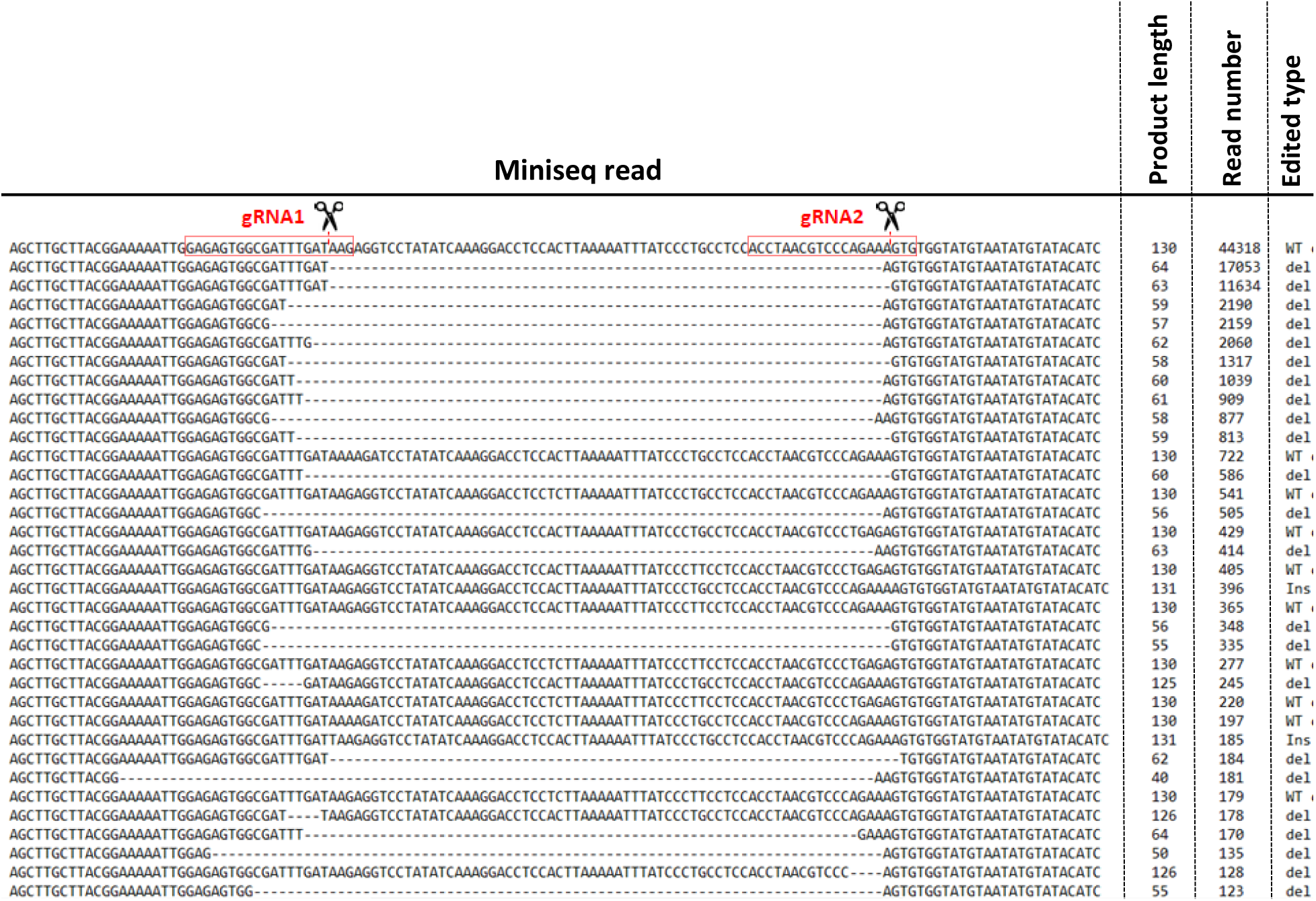
Indel alleles revealed from transfection of SpCas9, gR1.HPAT1 and gR2.HPAT3 with MJ.HPAT3-1 donor. The gRNA binding and cutting sites are shown on the top panel with the scissor icons on the WT sequence.

**Supplemental Figure 5.**
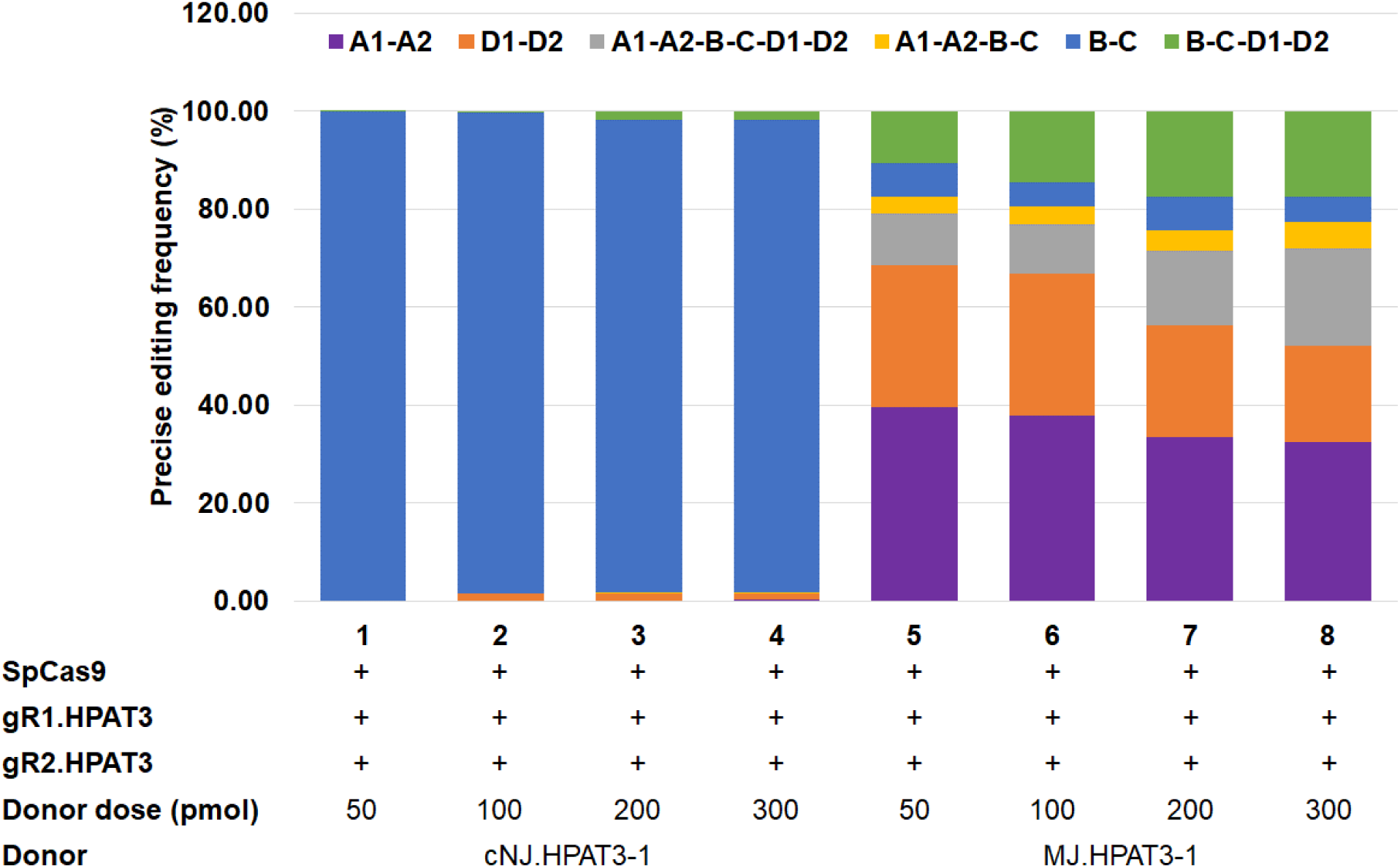
The frequency of the MMEJ-mediated repaired products at different donor doses.

**Supplemental Figure 6.**
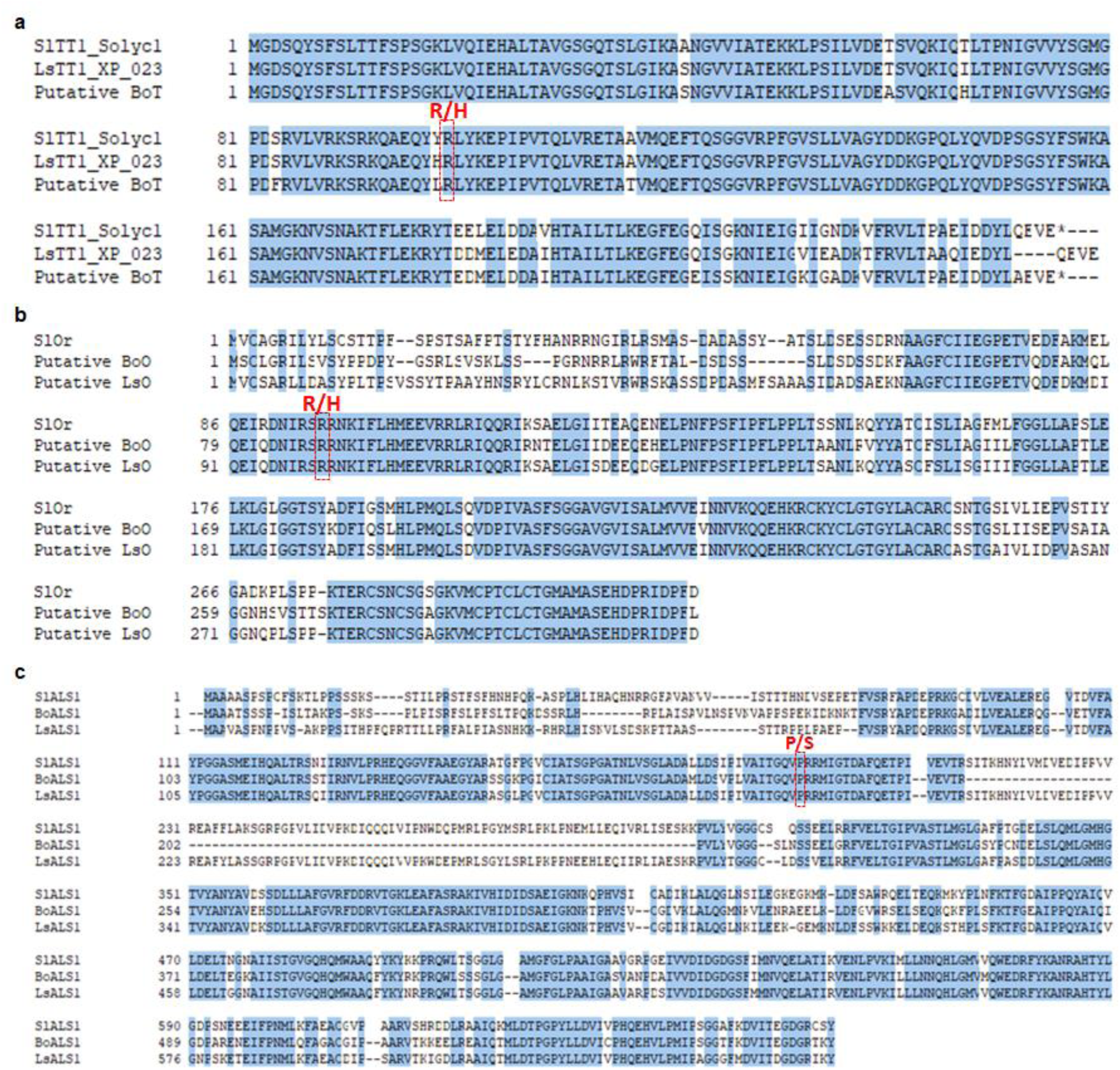
Identification of *TT1*, *Or*, and *ALS1* genes in lettuce and cabbage for MMEJ-mediated gene targeting. **a**-**c**, Protein sequence alignment among tomato, lettuce, and cabbage for identification of targeted a.a changes in *TT1* (**a**), *Or* (**b**), and *ALS1* (b) genes. The targeted a.a is highlighted by the blue discontinuous boxes.

**Supplemental Figure 7.**
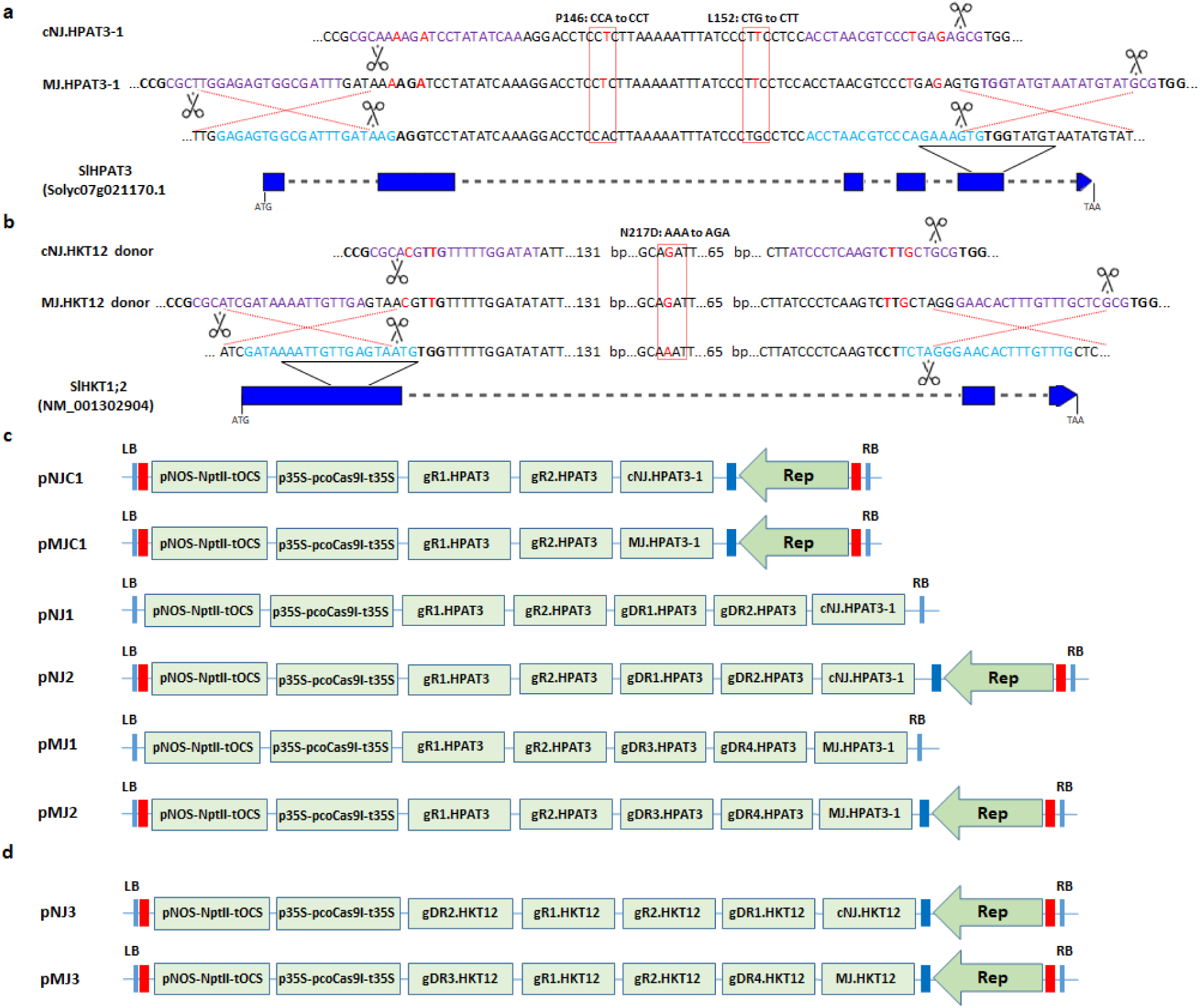
Agrobacterium-mediated system for cNHEJ and MMEJ-mediated gene editing in tomato. **a-b**, Schematic diagrams of cNHEJ- and MMEJ-mediated gene editing of tomato targeted loci (*SlHPAT3* (**a**) and *SlHKT1;2* (**b**) with their GenBank accession number), targeted sequences, gRNA binding sites, and donors. The targeted genes are shown from the start codon (ATG) to stop codons. Exons are drawn as blue boxes and the discontinuous lines in each gene represent intron parts of the gene; the gRNA binding sequences are highlighted in light blue color at the genomic sites and purple color at the donor vectors with PAMs (bold nucleotides). The expected cutting sites (3-nt upstream of a PAM) are indicated with the black scissors; the intended base changes are painted in the red font; the targeted amino acid changes and corresponding triplexes are denoted by the discontinuous red boxes and the texts placed above them. Two to three silent base changes corresponding to either the PAM or seed sequences (within 9 bases upstream of the 5’- NGG-3’ PAMs) were introduced into the donor to prevent the cleavage of the donor DNAs after they were successfully integrated into the genomic sites. The diagrams are not drawn to scale. **c-d**, Binary vectors used for tomato transformation to edit the *SlHPAT3* gene (**c**) and *SlHKT1;2* (**d**). A plant codon-optimized SpCas9 was employed to make the DSBs at the genomic sites and to release the donors. Also, a neomycin phosphotransferase II expression cassette was used for plant selection. The *SlHPAT3* gene was tested with control vectors (pNJC1 and pMJC1) without the donor cutting gRNAs. The pNJ1 and pMJ1 vectors are conventional T-DNAs and the other entire vectors are designed with a geminiviral replicon-based system that contains a Replicative protein expression cassette (Rep) driven by a bidirectional promoter (red bar, named as LIR) and terminated by a bidirectional terminator (blue bar, named as SIR). The replicon requires two LIRs to be autonomously replicated within the nuclei of plant cells.

**Supplemental Figure. 8.**
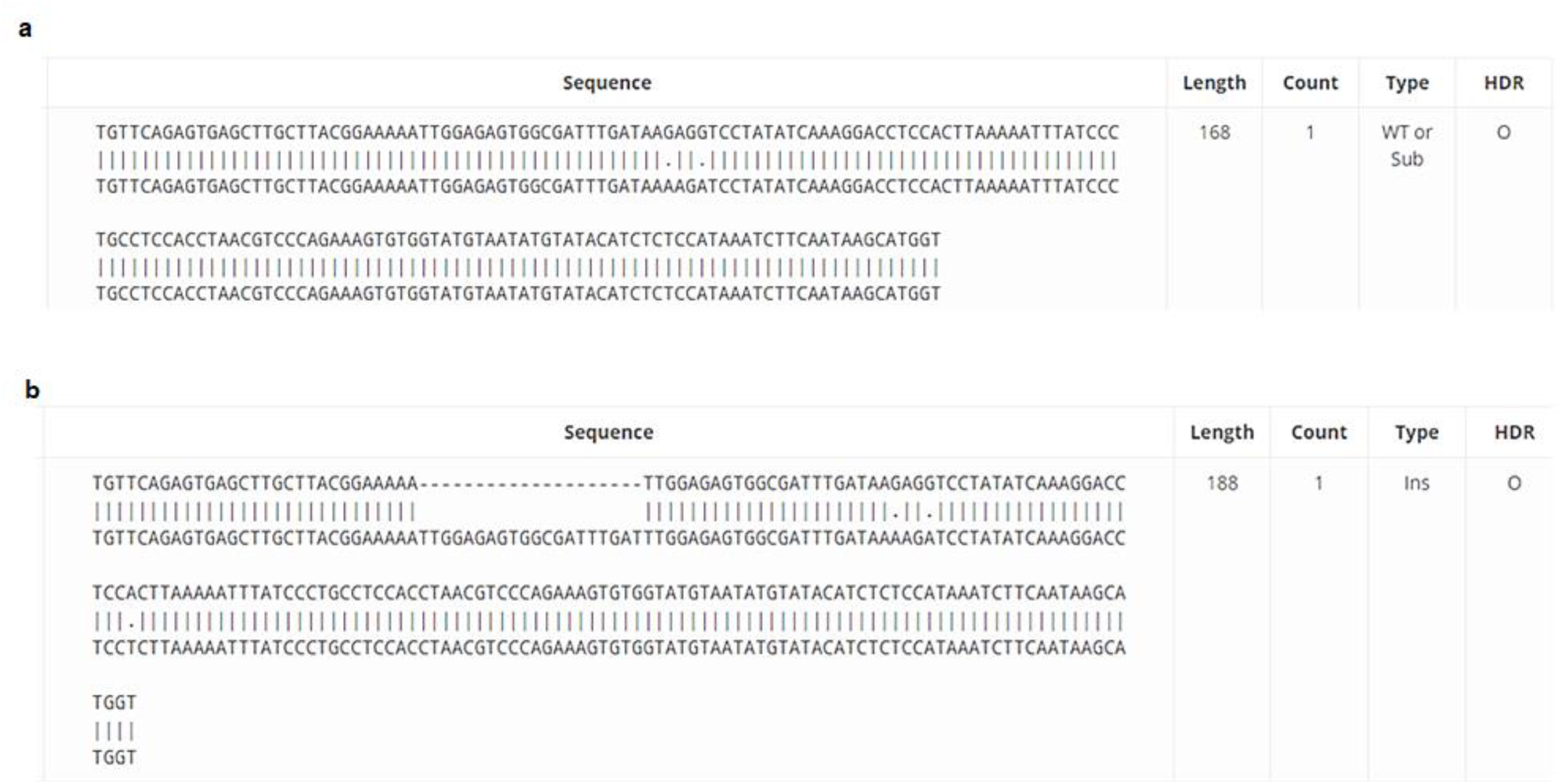
Representative repaired products obtained by the pMJ1 and pMJ2 vector. **a**, A one-sided precise replacement of the A1 and A2 bases by the pMJ1. **b**, A repaired product of the pMJ2 containing one-sided precise replacement of the A1 and A2 bases and insertion of the 20-bp 5’-microhomology arm.

**Supplemental Figure 9.**
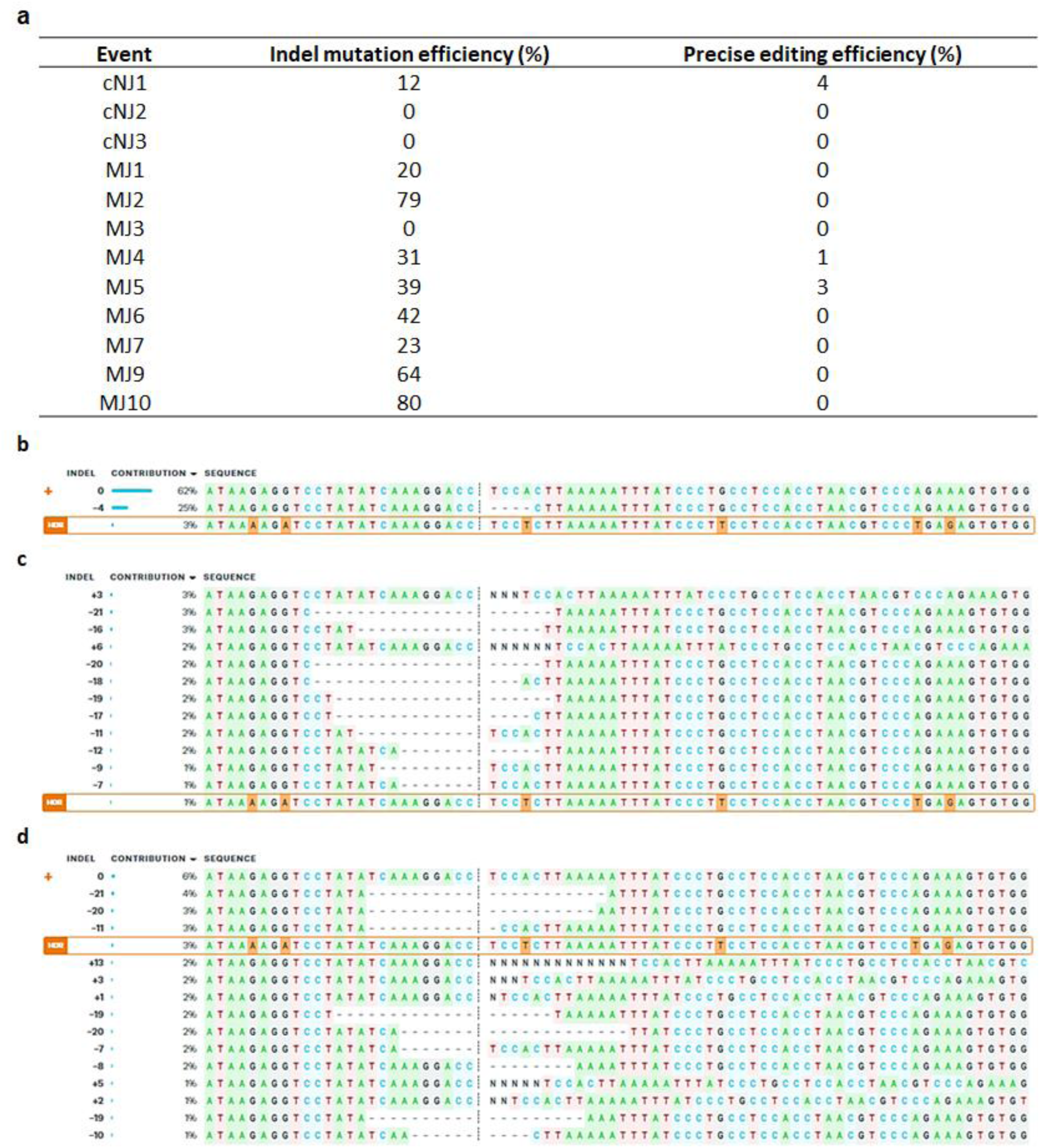
cNHEJ and MMEJ-mediated precision editing events revealed from *Agrobacterium*-mediated transformation. **a**. Table showing precisely edited allele frequency from regenerated plants transformed with the cNHEJ and MMEJ editing tools for editing the *SlHPAT3* locus. **b-d**. Edited alleles reported by ICE Synthego analysis with the cNHEJ event cNJ1 (**b**); the MMEJ events MJ4 (**c**) and MJ5 (**d**). The precisely edited alleles and efficiencies are denoted by the framed orange boxes named HDR that also highlighted modified bases in orange. The WT allele is marked with the plus sign.

## SUPPLEMENTAL TABLES

**Supplemental Table 1.**
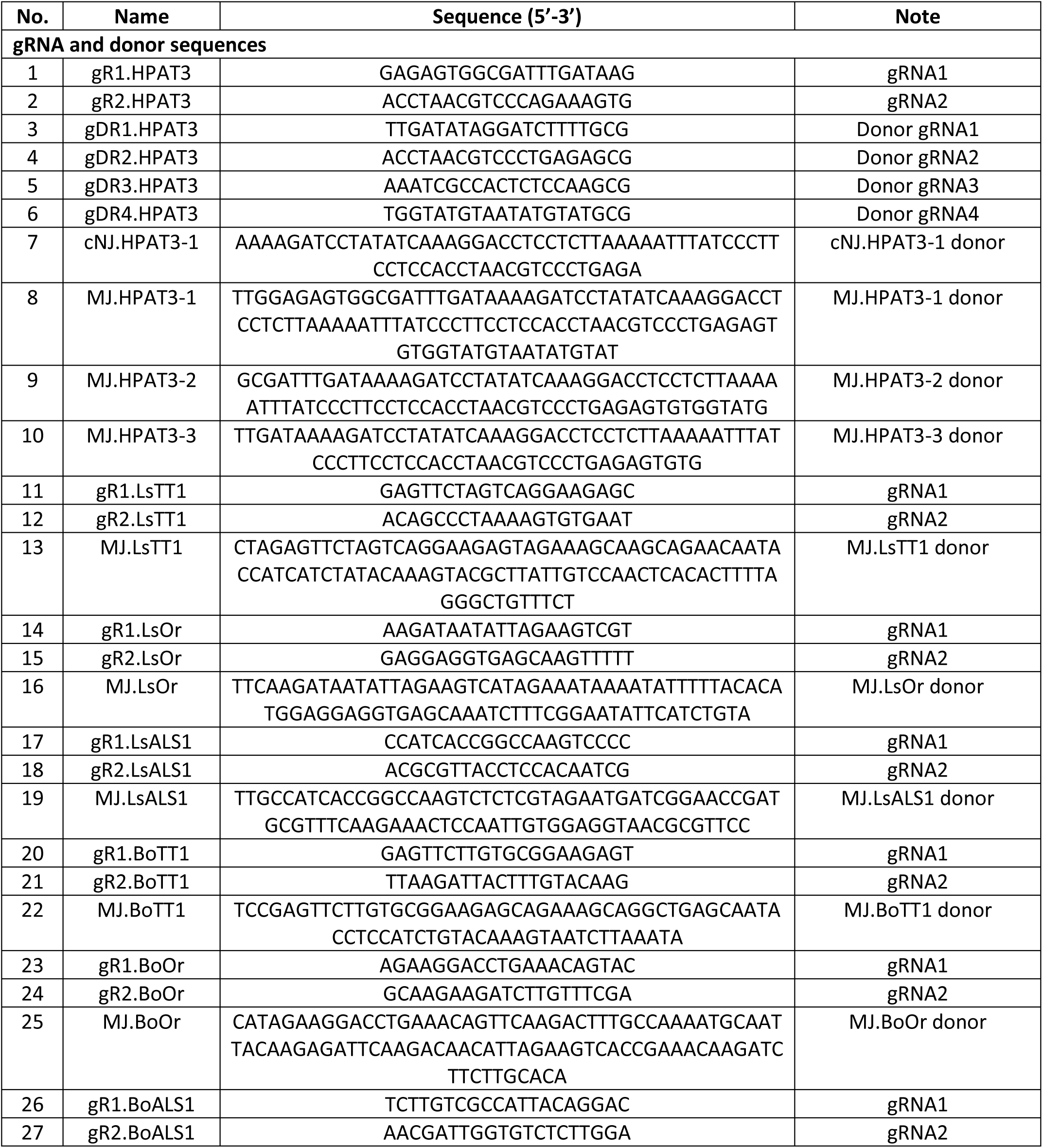
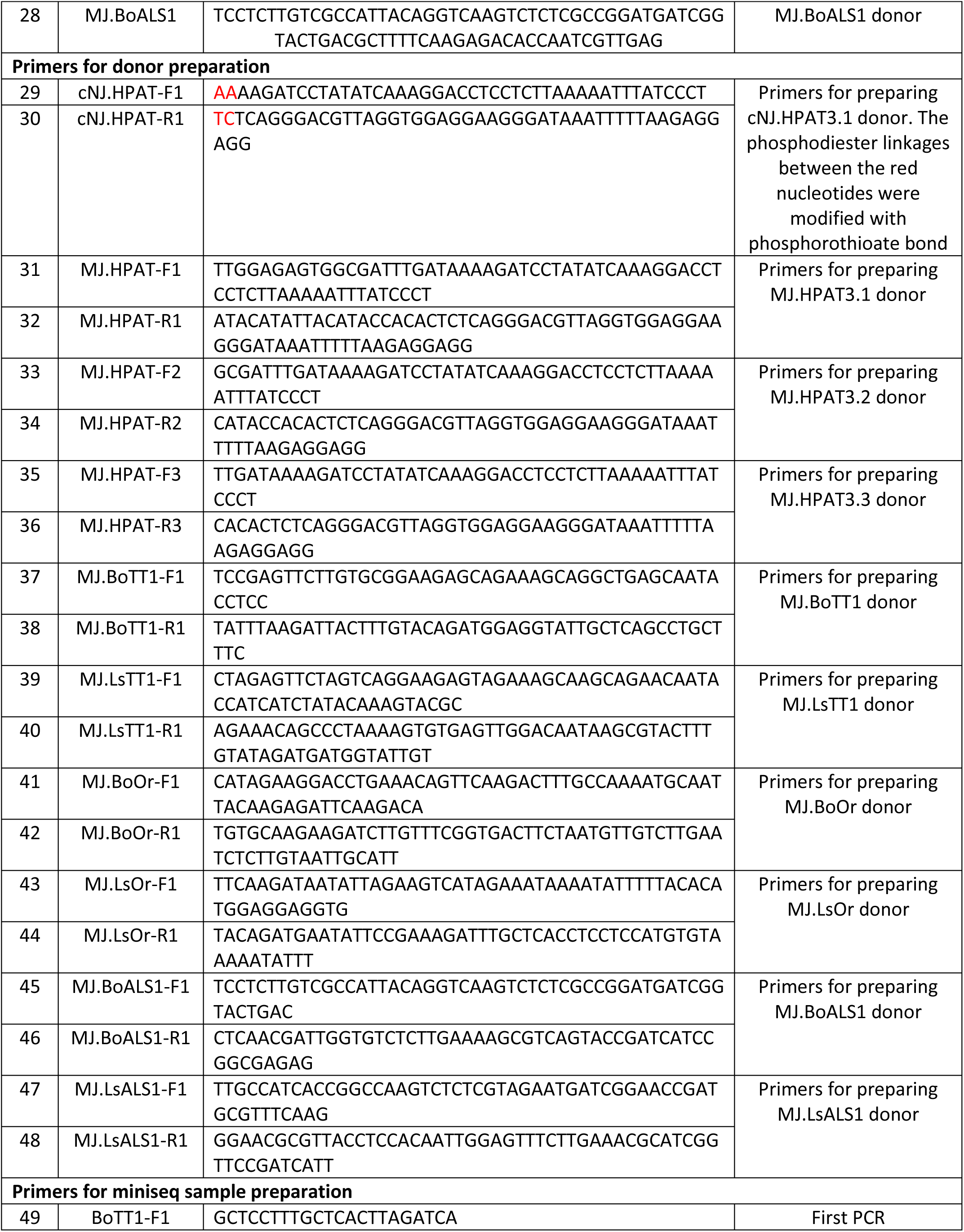
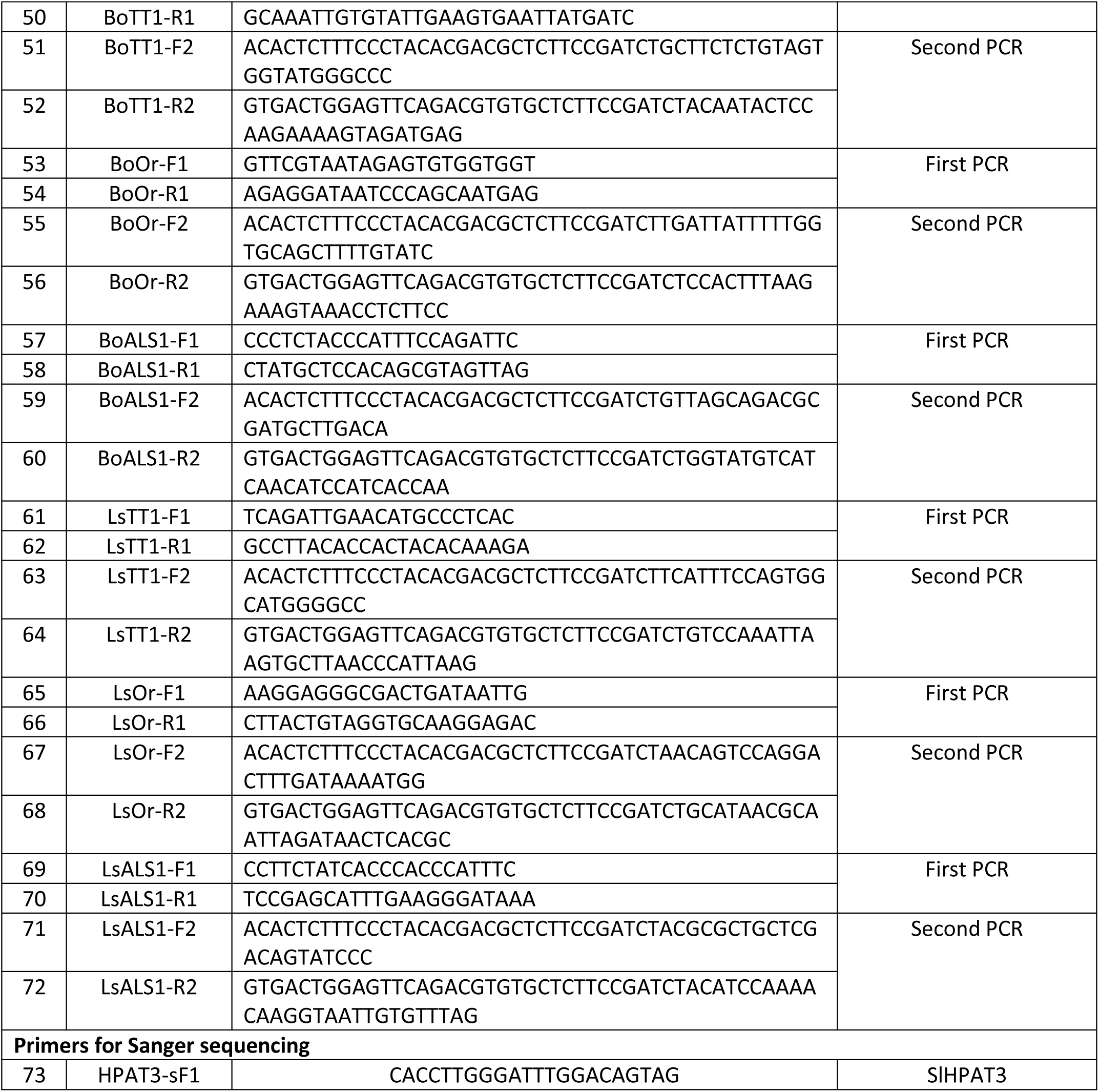
gRNA, donor, and primer sequences employed in the study.

**Supplemental Table 2.**
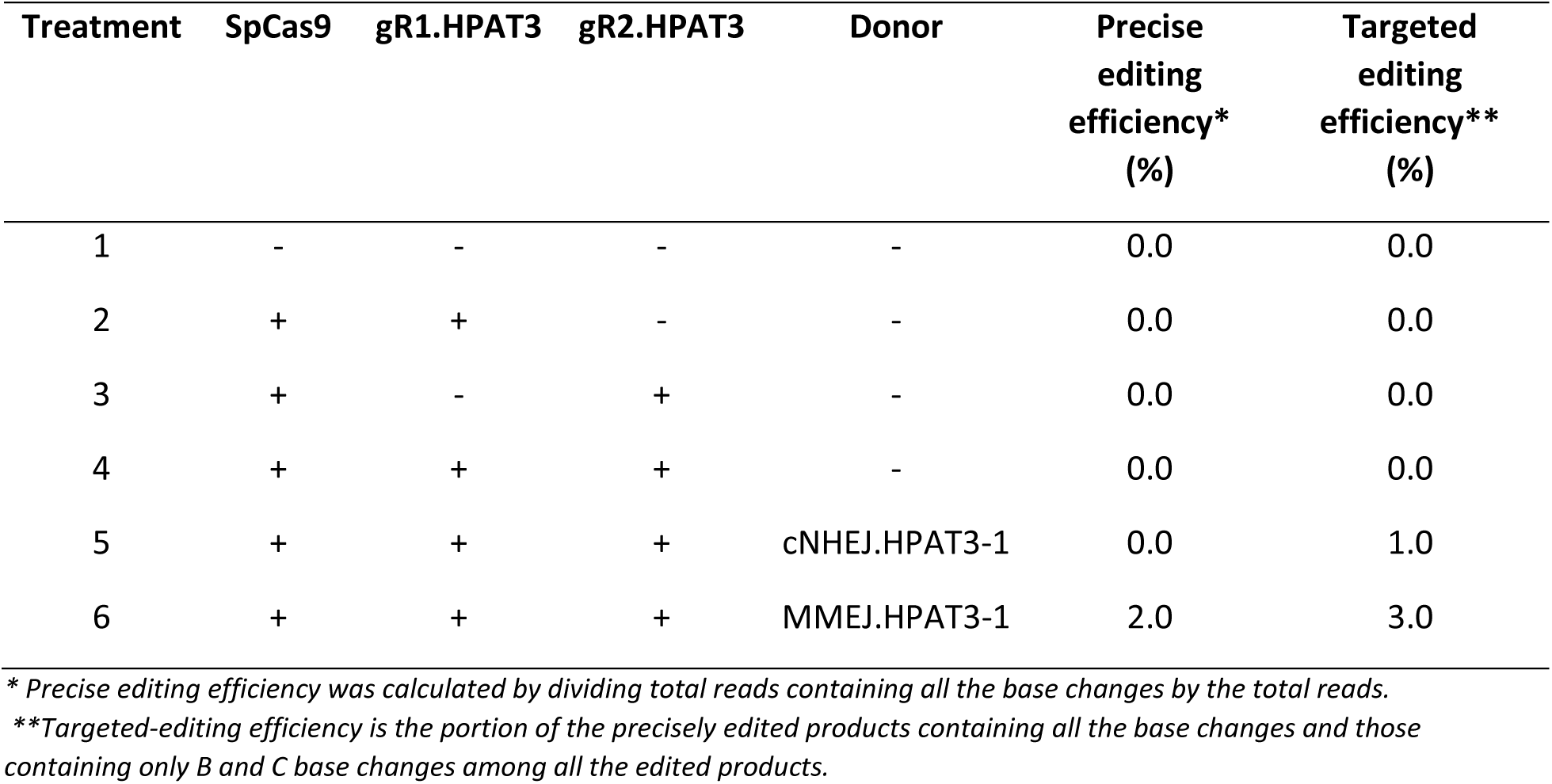
cNHEJ- and MMEJ-mediated editing rates at *SlHPAT3* locus revealed by Sanger sequencing.

**Supplemental Table 3.**
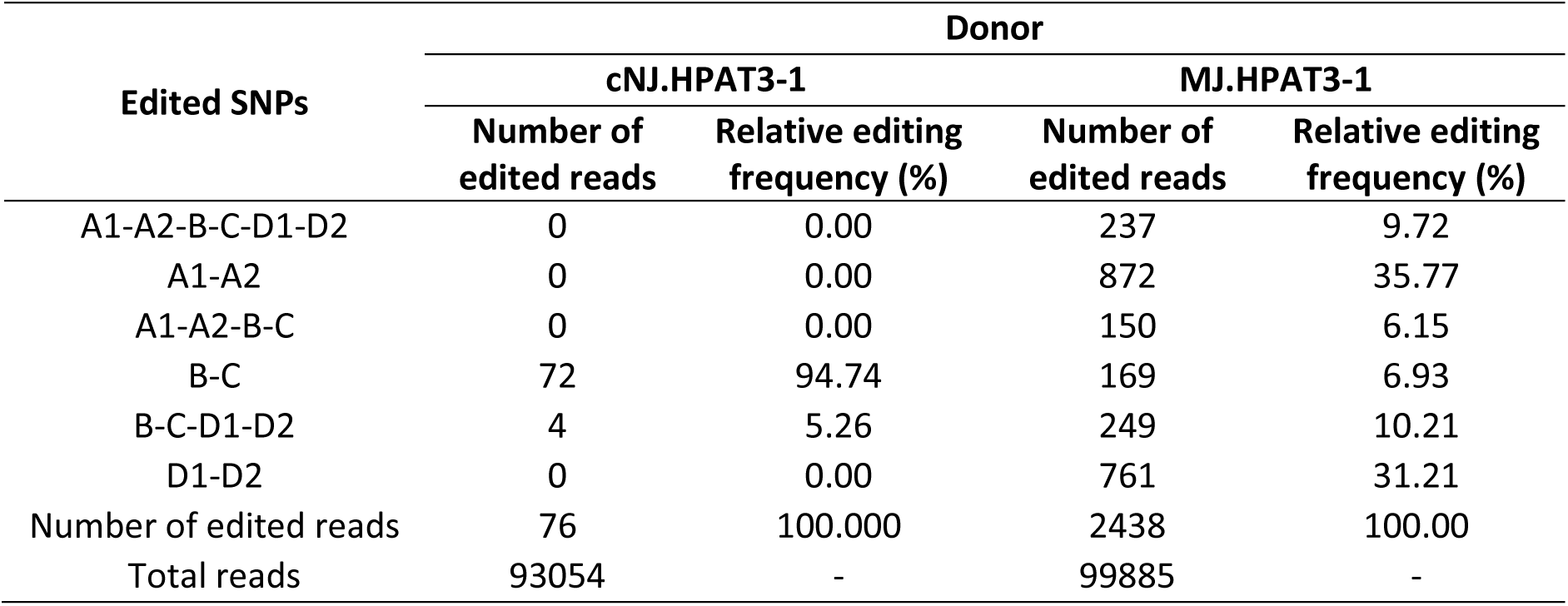
cNHEJ- and MMEJ-mediated editing rates of various repaired products at *SlHPAT3* locus.

**Supplemental Table 4.**
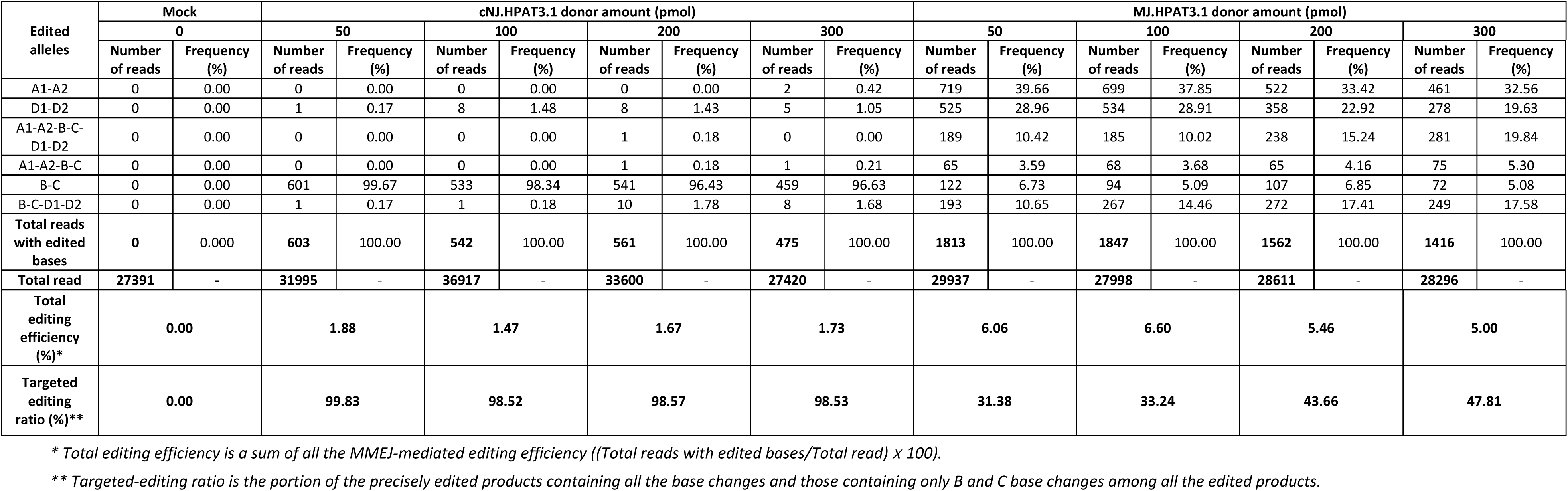
Editing frequency of various repaired products revealed from treatments of different donor types and doses.

**Supplemental Table 5.**
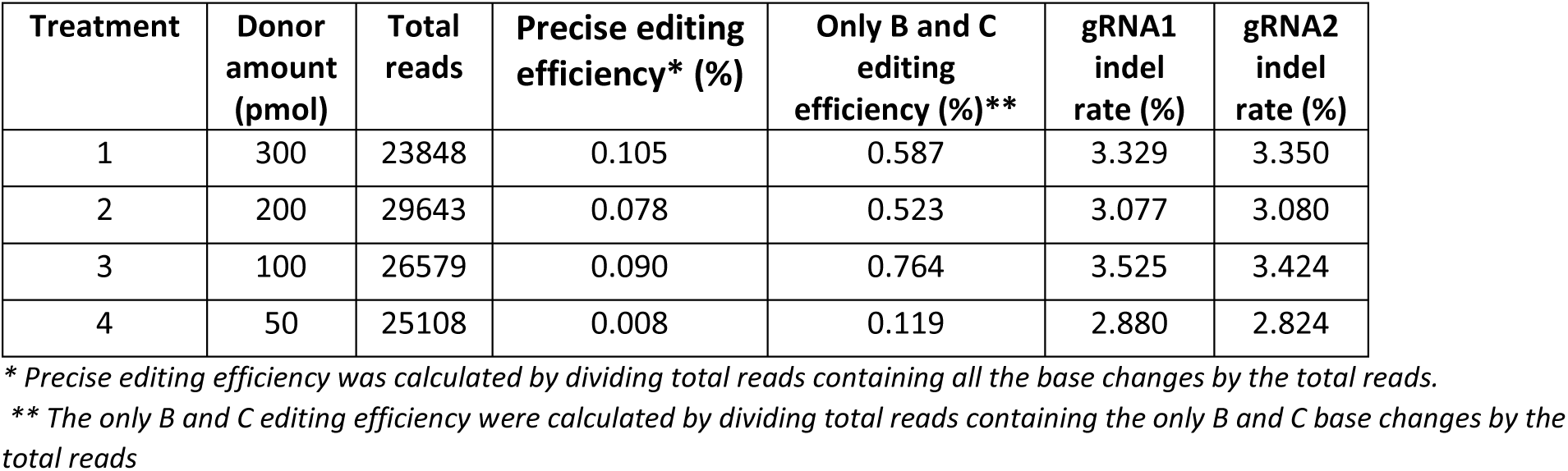
MMEJ-mediated gene editing using various MJ.HPAT3-1 donor amounts (replicate)

**Supplemental Table 6.**
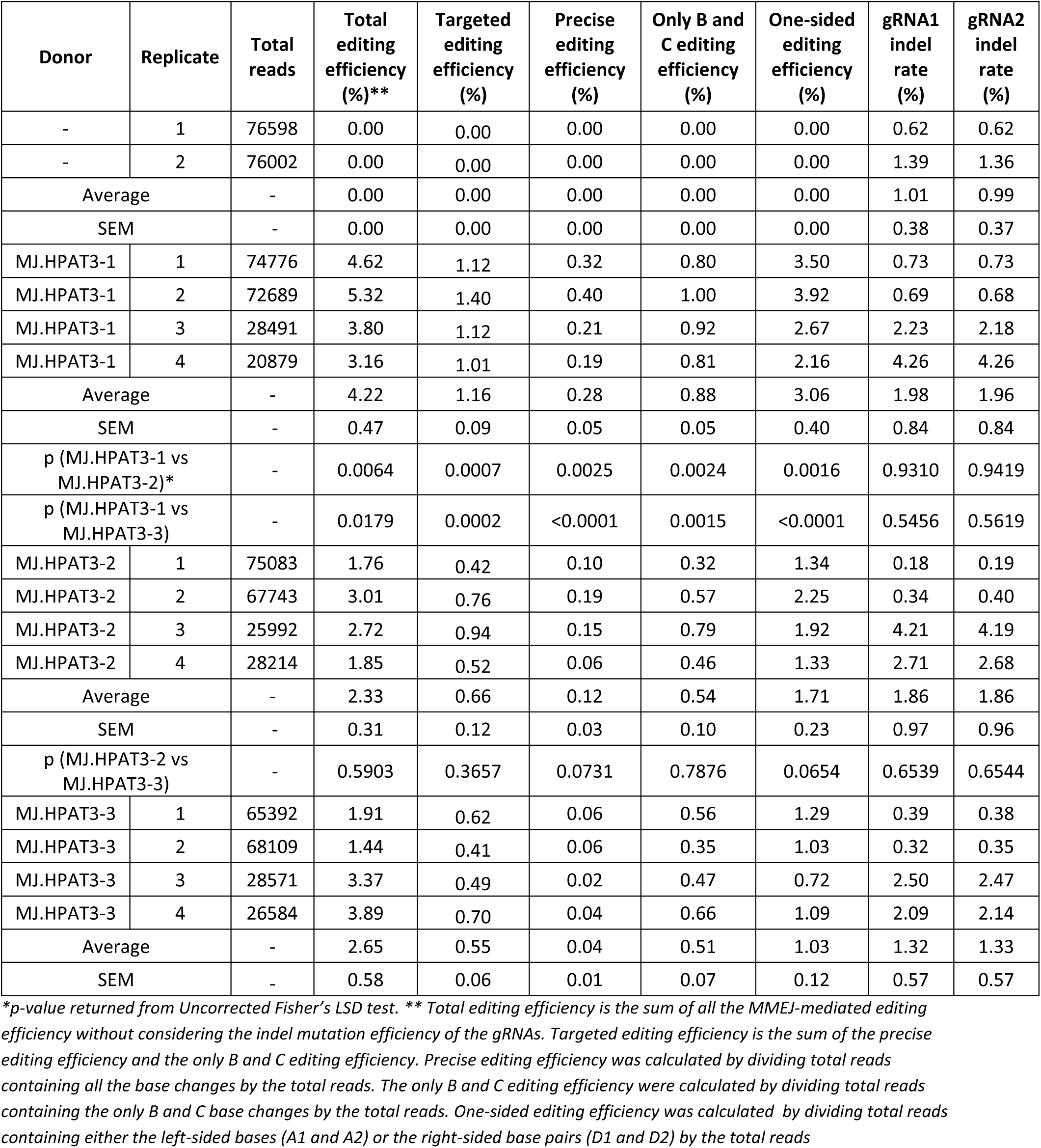
MMEJ-mediated editing efficiency revealed from DSBs repair using donors with different microhomology lengths.

**Supplemental Table 7.**
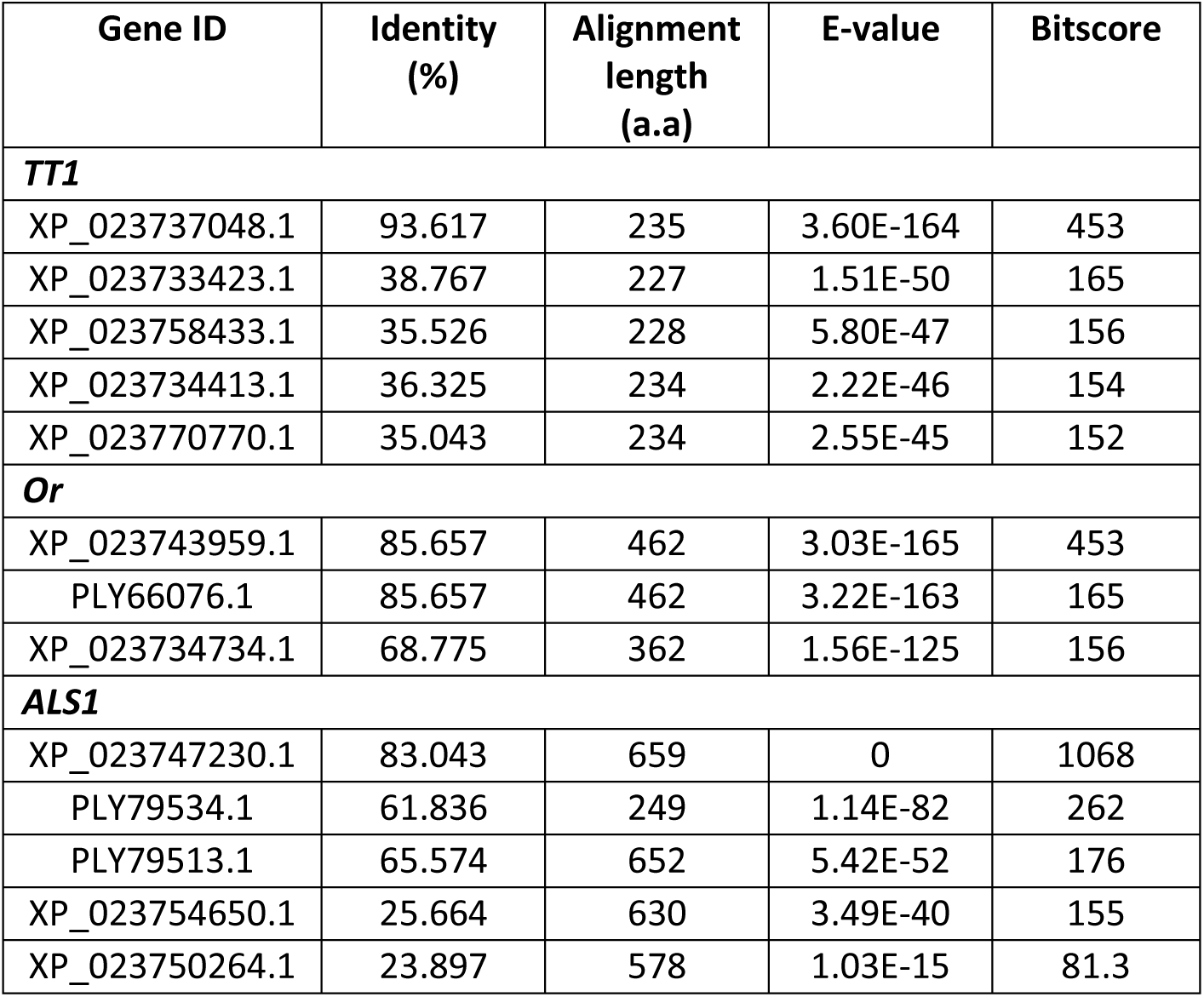
Sequence alignment for selection of targeted genes in lettuce.

**Supplemental Table 8.**
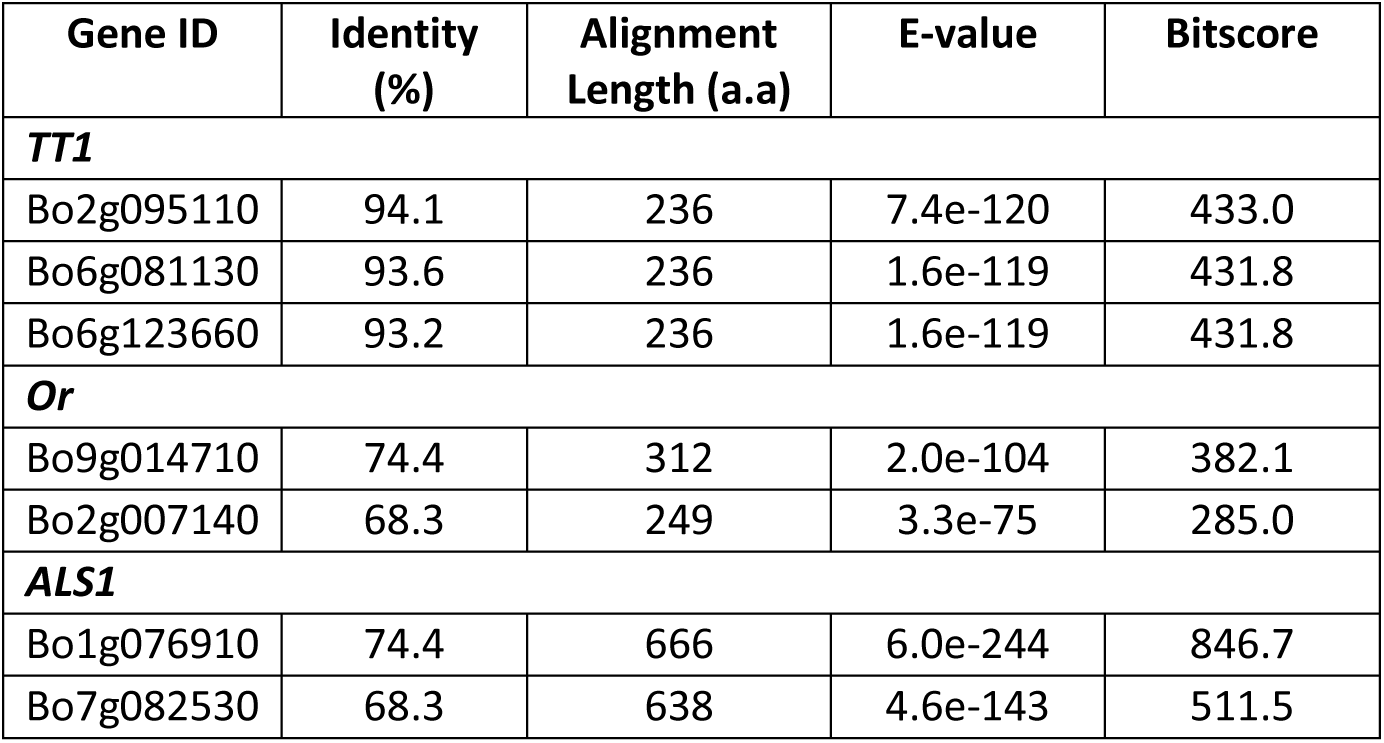
Sequence alignment for selection of targeted genes in cabbage.

**Supplemental Table 9.**
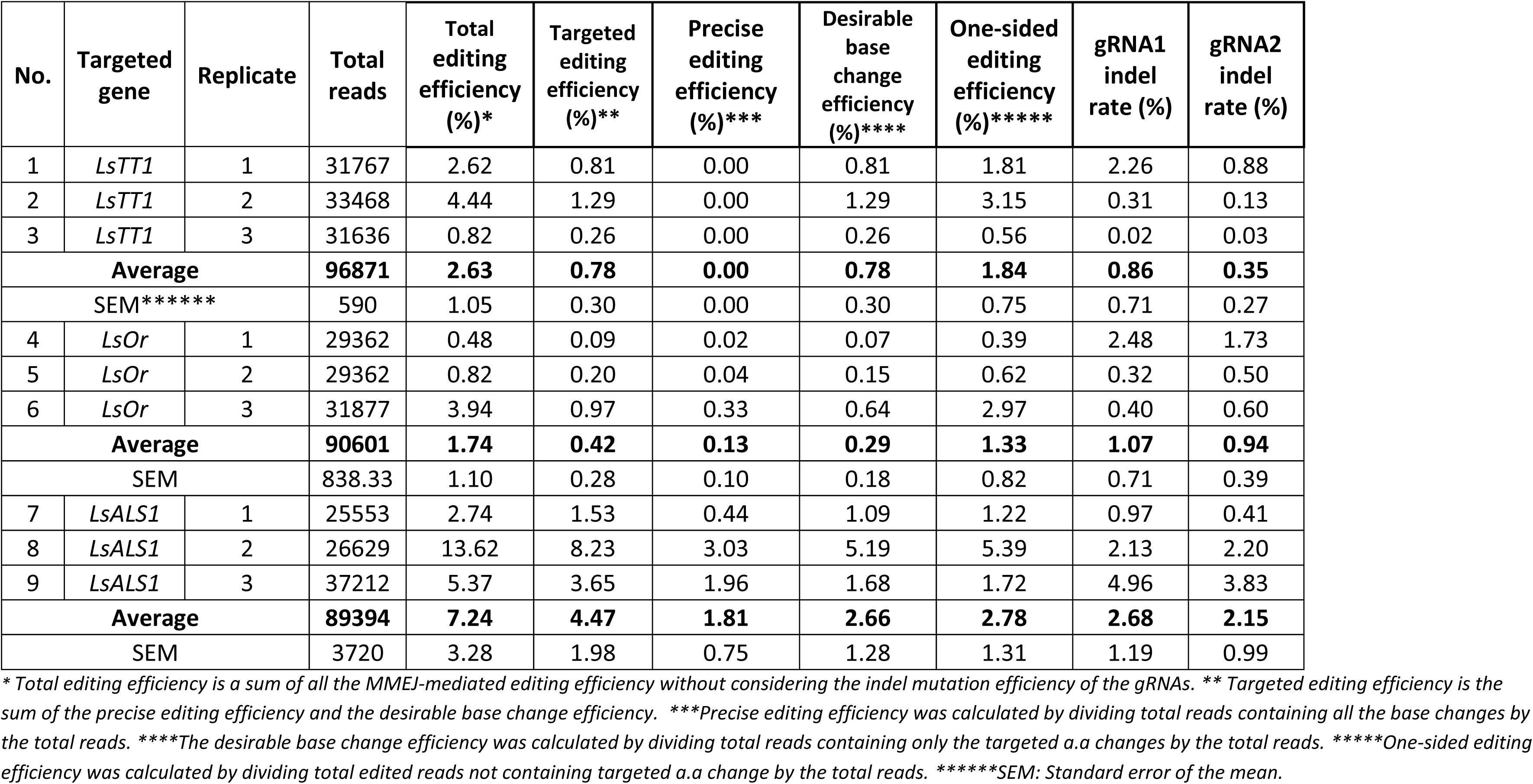
Data revealed from the targeted deep-sequencing analysis of MMEJ-mediated editing in lettuce.

**Supplemental Table 10.**
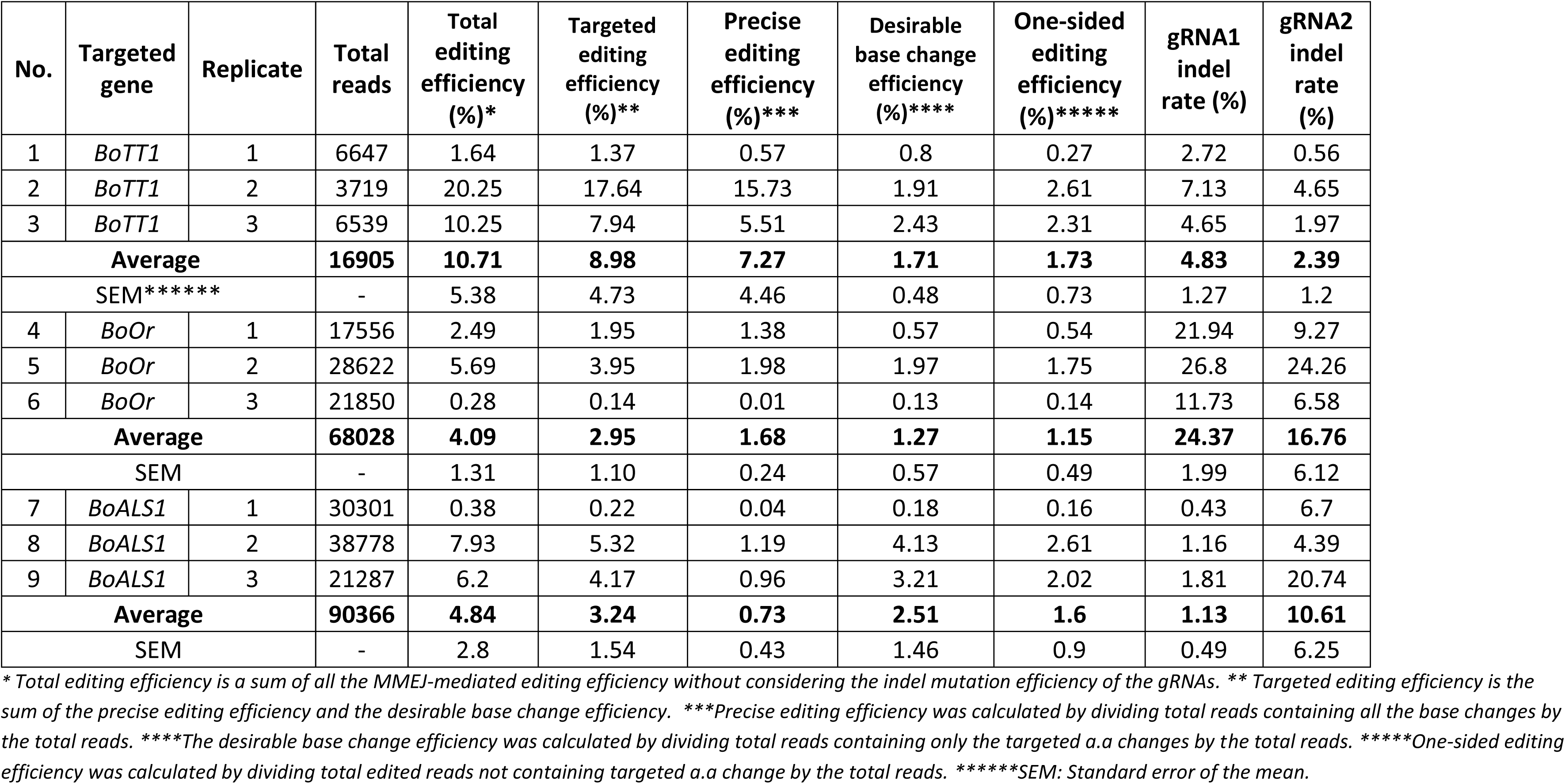
Data revealed from the targeted deep-sequencing analysis of MMEJ-mediated editing in cabbage.

**Supplementary Table 11.**
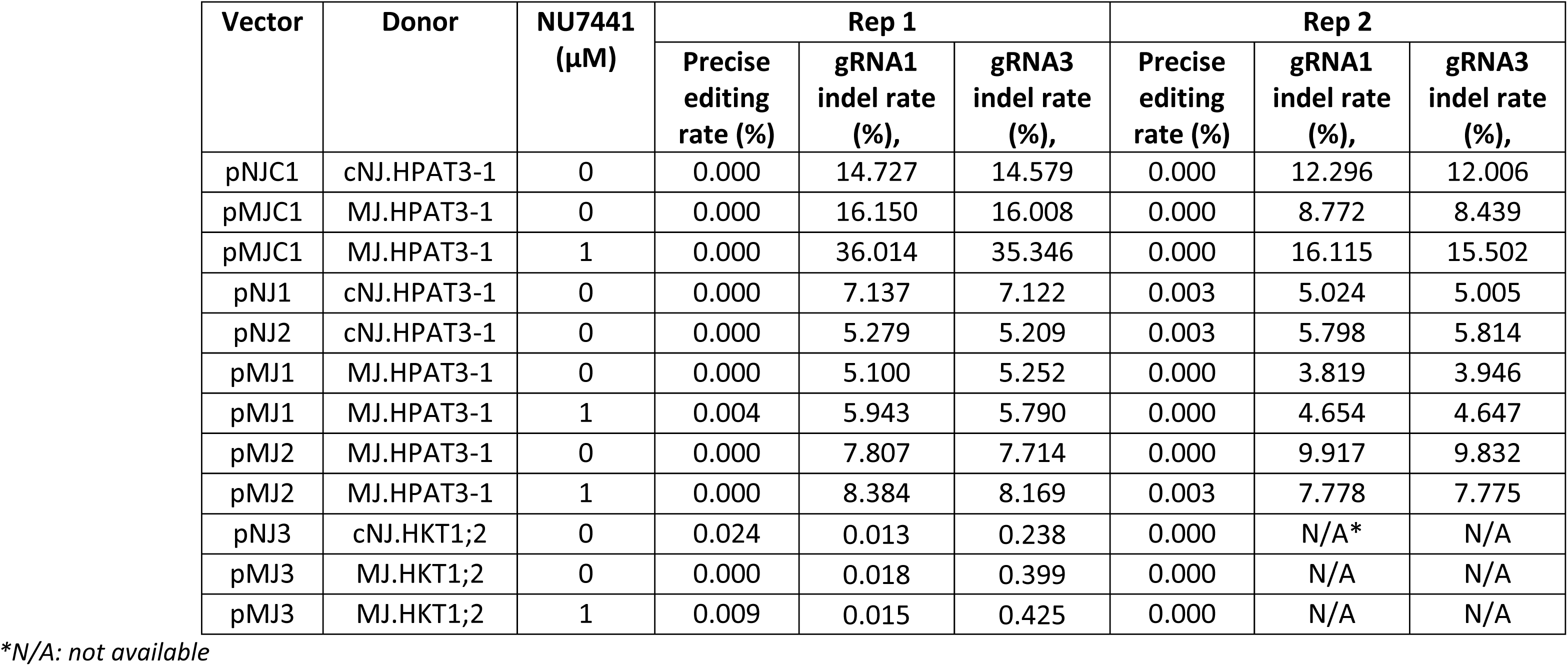
PREMJ-mediated gene editing in tomato using the *Agrobacterium*-mediated delivery of the editing tools.

## SEQUENCES USED IN THE STUDY

❖ **HPAT3 cNHEJ donor (cNJ.HPAT3-1) for RNP works** 5’-AAaAGaTCCTATATCAAAGGACCTCCtCTTAAAAATTTATCCCTtCCTCCACCTAACGTCCCtGAgA-3’ Blue font: introduced bases. From 5’-3’: the bases were name as A1; A2; B; C; D1; D2, respectively.
❖ **HPAT3 MMEJ donor (MJ.HPAT3-1) for RNP works** 5’- ATTGGAGAGTGGCGATTTGATAAaAGaTCCTATATCAAAGGACCTCCtCTTAAAAATTTATCCCTtCCTCCACCT AACGTCCCtGAgAGTGTGGTATGTAATATGTAT-3’ black font: replaced sequence; Blue font: introduced bases. From 5’-3’: the bases were name as A1; A2; B; C; D1; D2, respectively; red font: MH1; green font: MH2
❖ **HPAT3 MMEJ donor (MJ.HPAT3-2) for RNP works** 5’- GGCGATTTGATAAaAGaTCCTATATCAAAGGACCTCCtCTTAAAAATTTATCCCTtCCTCCACCTAACGTCCCtGA gAGTGTGGTATG-3’ black font: replaced sequence; Blue font: introduced bases. From 5’-3’: the bases were name as A1; A2; B; C; D1; D2, respectively; red font: MH1; green font: MH2
❖ **HPAT3 MMEJ donor (MJ.HPAT3-3) for RNP works** 5’- TTGATAAaAGaTCCTATATCAAAGGACCTCCtCTTAAAAATTTATCCCTtCCTCCACCTAACGTCCCtGAgAGTGT G-3’ black font: replaced sequence; Blue font: introduced bases. From 5’-3’: the bases were name as A1; A2; B; C; D1; D2, respectively; red font: MH1; green font: MH2
❖ **HPAT3 cNHEJ donor (cNJ.HPAT3-1) for *Agrobacterium*-mediated transformation works** CCgcgcAAaAGaTCCTATATCAAAGGACCTCCtCTTAAAAATTTATCCCTtCCTCCACCTAACGTCCCtGAgAgcgtG G Blue font: introduced bases. From 5’-3’: the bases were name as A1; A2; B; C; D1; D2, respectively; yellow highlighted font: gDR1.HPAT3 binding sequence; gray highlighted font: gDR2.HPAT3 binding sequence.
❖ **HPAT3 MMEJ donor (MJ.HPAT3-1) for *Agrobacterium*-mediated transformation works** CCgcgcTTGGAGAGTGGCGATTTGATAAaAGaTCCTATATCAAAGGACCTCCtCTTAAAAATTTATCCCTtCCTCC ACCTAACGTCCCtGAgAGTGTGGTATGTAATATGTATgcgtGG Blue font: introduced bases. From 5’-3’: the bases were name as A1; A2; B; C; D1; D2, respectively; yellow highlighted font: gDR3.HPAT3 binding sequence; gray highlighted font: gDR4.HPAT3 binding sequence. red font: MH1; green font: MH2
❖ **sgR1.HPAT3** GAGAGTGGCGATTTGATAAGgttttagagctagaaatagcaagttaaaataaggctagtccgttatcaacttgaaaaagtg gcaccgagtcggtgc Black font: gRNA sequence; green font: CRISPR gRNA scaffold.
❖ **sgR2.HPAT3** ACCTAACGTCCCAGAAAGTGgttttagagctagaaatagcaagttaaaataaggctagtccgttatcaacttgaaaaagtgg caccgagtcggtgc Black font: gRNA sequence; green font: CRISPR gRNA scaffold.
❖ **gDR1.HPAT3** TTGATATAGGAtCTtTTgcggttttagagctagaaatagcaagttaaaataaggctagtccgttatcaacttgaaaaagtggcaccgagtc ggtgcTTTTTTT Black font: gRNA sequence; green font: CRISPR gRNA scaffold.
❖ **gDR2.HPAT3** ACCTAACGTCCCtGAgAgcggttttagagctagaaatagcaagttaaaataaggctagtccgttatcaacttgaaaaagtggcaccgagt cggtgcTTTTTTT Black font: gRNA sequence; green font: CRISPR gRNA scaffold.
❖ **gDR3.HPAT3** AAATCGCCACTCTCCAAgcggttttagagctagaaatagcaagttaaaataaggctagtccgttatcaacttgaaaaagtggcaccgagt cggtgcTTTTTTT Black font: gRNA sequence; green font: CRISPR gRNA scaffold.
❖ **gDR4.HPAT3** TGGTATGTAATATGTATgcggttttagagctagaaatagcaagttaaaataaggctagtccgttatcaacttgaaaaagtggcaccgag tcggtgcTTTTTTT Black font: gRNA sequence; green font: CRISPR gRNA scaffold.
❖ **HKT1;2 cNHEJ donor (cNJ.HKT12) for *Agrobacterium*-mediated transformation works** CCgcgcAcGTtGTTTTTGGATATATTCTTGTTGTTATTCTTCTTGGTTCATCGTTAGTTTCTCTCTATATAATAATCA TCCCTAGCGCCAAACAAATCCTTGACCAAAAAGGCCTTAATTTACATACTTTTTCACTATTCACCACAGTATCAA CTTTTGCAgATTGTGGTTTTTTACCTACAAATGAAAACATGATGATTTTCAAGAAAAATTCAGGaCTTCTTCTCA TTCTTATCCCTCAAGTCtTgCTgcgtGG Blue font: introduced bases; yellow highlighted font: gDR1.HKT12 binding sequence; gray highlighted font: gDR2.HKT12 binding sequence.
❖ **HKT1;2 MMEJ donor (MJ.HKT12) for *Agrobacterium*-mediated transformation works** CCgcgcATCGATAAAATTGTTGAGTAAcGTtGTTTTTGGATATATTCTTGTTGTTATTCTTCTTGGTTCATCGTTAG TTTCTCTCTATATAATAATCATCCCTAGCGCCAAACAAATCCTTGACCAAAAAGGCCTTAATTTACATACTTTTT CACTATTCACCACAGTATCAACTTTTGCAgATTGTGGTTTTTTACCTACAAATGAAAACATGATGATTTTCAAGA AAAATTCAGGaCTTCTTCTCATTCTTATCCCTCAAGTCtTgCTAGGGAACACTTTGTTTGCTCgcgtGG Blue font: introduced bases. yellow highlighted font: gDR3.HKT12 binding sequence; gray highlighted font: gDR4.HKT12 binding sequence. red font: MH1; green font: MH2
❖ **sgR1.HKT12** GATAAAATTGTTGAGTAATGgttttagagctagaaatagcaagttaaaataaggctagtccgttatcaacttgaaaaagtggcaccga gtcggtgcTTTTTTT Black font: gRNA sequence; green font: CRISPR gRNA scaffold.
❖ **sgR2.HKT12** CAAACAAAGTGTTCCCTAGAgttttagagctagaaatagcaagttaaaataaggctagtccgttatcaacttgaaaaagtggcaccga gtcggtgcTTTTTTT Black font: gRNA sequence; green font: CRISPR gRNA scaffold.
❖ **gDR1.HKT12** ATATCCAAAAACaACgTgcggttttagagctagaaatagcaagttaaaataaggctagtccgttatcaacttgaaaaagtggcaccgagt cggtgcTTTTTTT Black font: gRNA sequence; green font: CRISPR gRNA scaffold.
❖ **gDR2.HKT12** ATCCCTCAAGTCtTgCTgcggttttagagctagaaatagcaagttaaaataaggctagtccgttatcaacttgaaaaagtggcaccgagtc ggtgcTTTTTTT Black font: gRNA sequence; green font: CRISPR gRNA scaffold.
❖ **gDR3.HKT12** TCAACAATTTTATCGATgcggttttagagctagaaatagcaagttaaaataaggctagtccgttatcaacttgaaaaagtggcaccgagt cggtgcTTTTTTT Black font: gRNA sequence; green font: CRISPR gRNA scaffold.
❖ **gDR4.HKT12** GAACACTTTGTTTGCTCgcggttttagagctagaaatagcaagttaaaataaggctagtccgttatcaacttgaaaaagtggcaccgagt cggtgcTTTTT Black font: gRNA sequence; green font: CRISPR gRNA scaffold.
❖ p35S-pcoCas9I-t35S sequence GGAGGAATTCCAATCCCACAAAAATCTGAGCTTAACAGCACAGTTGCTCCTCTCAGAGCAGAATCGGGTATTCAAC ACCCTCATATCAACTACTACGTTGTGTATAACGGTCCACATGCCGGTATATACGATGACTGGGGTTGTACAAAGGC GGCAACAAACGGCGTTCCCGGAGTTGCACACAAGAAATTTGCCACTATTACAGAGGCAAGAGCAGCAGCTGACGC GTACACAACAAGTCAGCAAACAGACAGGTTGAACTTCATCCCCAAAGGAGAAGCTCAACTCAAGCCCAAGAGCTT TGCTAAGGCCCTAACAAGCCCACCAAAGCAAAAAGCCCACTGGCTCACGCTAGGAACCAAAAGGCCCAGCAGTGA TCCAGCCCCAAAAGAGATCTCCTTTGCCCCGGAGATTACAATGGACGATTTCCTCTATCTTTACGATCTAGGAAGG AAGTTCGAAGGTGAAGGTGACGACACTATGTTCACCACTGATAATGAGAAGGTTAGCCTCTTCAATTTCAGAAAG AATGCTGACCCACAGATGGTTAGAGAGGCCTACGCAGCAAGTCTCATCAAGACGATCTACCCGAGTAACAATCTCC AGGAGATCAAATACCTTCCCAAGAAGGTTAAAGATGCAGTCAAAAGATTCAGGACTAATTGCATCAAGAACACAG AGAAAGACATATTTCTCAAGATCAGAAGTACTATTCCAGTATGGACGATTCAAGGCTTGCTTCATAAACCAAGGCA AGTAATAGAGATTGGAGTCTCTAAAAAGGTAGTTCCTACTGAATCTAAGGCCATGCATGGAGTCTAAGATTCAAAT CGAGGATCTAACAGAACTCGCCGTCAAGACTGGCGAACAGTTCATACAGAGTCTTTTACGACTCAATGACAAGAA GAAAATCTTCGTCAACATGGTGGAGCACGACACTCTGGTCTACTCCAAAAATGTCAAAGATACAGTCTCAGAAGAT CAAAGGGCTATTGAGACTTTTCAACAAAGGATAATTTCGGGAAACCTCCTCGGATTCCATTGCCCAGCTATCTGTC ACTTCATCGAAAGGACAGTAGAAAAGGAAGGTGGCTCCTACAAATGCCATCATTGCGATAAAGGAAAGGCTATCA TTCAAGATCTCTCTGCCGACAGTGGTCCCAAAGATGGACCCCCACCCACGAGGAGCATCGTGGAAAAAGAAGAGG TTCCAACCACGTCTACAAAGCAAGTGGATTGATGTGACATCTCCACTGACGTAAGGGATGACGCACAATCCCACTA TCCTTCGCAAGACCCTTCCTCTATATAAGGAAGTTCATTTCATTTGGAGAGGACACGCTCGAGTATAAGGTAAATTT CTGTGTTCCTTATTCTCTCAAAATCTTCGATTTTGTTTTCGTTCGATCCCAATTTCGTATATGTTCTTTGGTTTAGATTC TGTTAATCTTAGATCGAAGATGATTTTCTGGGTTTGATCGTTAGATATCATCTTAATTCTCGATTAGGGTTTCATAG ATATCATCCGATTTGTTCAAATAATTTGAGTTTTGTCGAATAATTACTCTTCGATTTGTGATTTCTATCTAGATCTGG TGTTAGTTTCTAGTTTGTGCGATCGAATTTGTCGATTAATCTGAGTTTTTCTGATTAACAGGAGCTCATTTTTACAAC AATTACCAACAACAACAAACAACAAACAACATTACAATTACATTTACAATTATCGATACAATGacc ATGAAACGGACAGCCGACGGAAGCGAGTTCGAGGCCCCAAAGAAGAAGCGGAAGGTCGATAAGAAGTACTCTAT CGGACTCGATATCGGAACTAACTCTGTGGGATGGGCTGTGATCACCGATGAGTACAAGGtaaagcctcgatttttgggtttaggtgtctgcttattagagtaaaaacacatcctttgaaattgtttgtggtcatttgattgtgctcttgatccattgaattgctGCAGGTGCCATCTAA GAAGTTCAAGGTTCTCGGAAACACCGATAGGCACTCTATCAAGAAAAACCTTATCGGTGCTCTCCTCTTCGATTCTG GTGAAACTGCTGAGGCTACCAGACTCAAGAGAACCGCTAGAAGAAGGTACACCAGAAGAAAGAACAGGATCTGC TACCTCCAAGAGATCTTCTCTAACGAGATGGCTAAAGTGGATGATTCATTCTTCCACAGGCTCGAAGAGTCATTCCT CGTGGAAGAAGATAAGAAGCACGAGAGGCACCCTATCTTCGGAAACATCGTTGATGAGGTGGCATACCACGAGA AGTACCCTACTATCTACCACCTCAGAAAGAAGCTCGTTGATTCTACTGATAAGGCTGATCTCAGGCTCATCTACCTC GCTCTCGCTCACATGATCAAGTTCAGAGGACACTTCCTCATCGAGGGTGATCTCAACCCTGATAACTCTGATGTGG ATAAGTTGTTCATCCAGCTCGTGCAGACCTACAACCAGCTTTTCGAAGAGAACCCTATCAACGCTTCAGGTGTGGA TGCTAAGGCTATCCTCTCTGCTAGGCTCTCTAAGTCAAGAAGGCTTGAGAACCTCATTGCTCAGCTCCCTGGTGAG AAGAAGAACGGACTTTTCGGAAACTTGATCGCTCTCTCTCTCGGACTCACCCCTAACTTCAAGTCTAACTTCGATCT CGCTGAGGATGCAAAGCTCCAGCTCTCAAAGGATACCTACGATGATGATCTCGATAACCTCCTCGCTCAGATCGGA GATCAGTACGCTGATTTGTTCCTCGCTGCTAAGAACCTCTCTGATGCTATCCTCCTCAGTGATATCCTCAGAGTGAA CACCGAGATCACCAAGGCTCCACTCTCAGCTTCTATGATCAAGAGATACGATGAGCACCACCAGGATCTCACACTT CTCAAGGCTCTTGTTAGACAGCAGCTCCCAGAGAAGTACAAAGAGATTTTCTTCGATCAGTCTAAGAACGGATACG CTGGTTACATCGATGGTGGTGCATCTCAAGAAGAGTTCTACAAGTTCATCAAGCCTATCCTCGAGAAGATGGATGG AACCGAGGAACTCCTCGTGAAGCTCAATAGAGAGGATCTTCTCAGAAAGCAGAGGACCTTCGATAACGGATCTAT CCCTCATCAGATCCACCTCGGAGAGTTGCACGCTATCCTTAGAAGGCAAGAGGATTTCTACCCATTCCTCAAGGAT AACAGGGAAAAGATTGAGAAGATTCTCACCTTCAGAATCCCTTACTACGTGGGACCTCTCGCTAGAGGAAACTCAA GATTCGCTTGGATGACCAGAAAGTCTGAGGAAACCATCACCCCTTGGAACTTCGAAGAGGTGGTGGATAAGGGT GCTAGTGCTCAGTCTTTCATCGAGAGGATGACCAACTTCGATAAGAACCTTCCAAACGAGAAGGTGCTCCCTAAGC ACTCTTTGCTCTACGAGTACTTCACCGTGTACAACGAGTTGACCAAGGTTAAGTACGTGACCGAGGGAATGAGGA AGCCTGCTTTTTTGTCAGGTGAGCAAAAGAAGGCTATCGTTGATCTCTTGTTCAAGACCAACAGAAAGGTGACCGT GAAGCAGCTCAAAGAGGATTACTTCAAGAAAATCGAGTGCTTCGATTCAGTTGAGATTTCTGGTGTTGAGGATAG GTTCAACGCATCTCTCGGAACCTACCACGATCTCCTCAAGATCATTAAGGATAAGGATTTCTTGGATAACGAGGAA AACGAGGATATCTTGGAGGATATCGTTCTTACCCTCACCCTCTTTGAAGATAGAGAGATGATTGAAGAAAGGCTCA AGACCTACGCTCATCTCTTCGATGATAAGGTGATGAAGCAGTTGAAGAGAAGAAGATACACTGGTTGGGGAAGG CTCTCAAGAAAGCTCATTAACGGAATCAGGGATAAGCAGTCTGGAAAGACAATCCTTGATTTCCTCAAGTCTGATG GATTCGCTAACAGAAACTTCATGCAGCTCATCCACGATGATTCTCTCACCTTTAAAGAGGATATCCAGAAGGCTCA GGTTTCAGGACAGGGTGATAGTCTCCATGAGCATATCGCTAACCTCGCTGGATCTCCTGCAATCAAGAAGGGAAT CCTCCAGACTGTGAAGGTTGTGGATGAGTTGGTGAAGGTGATGGGAAGGCATAAGCCTGAGAACATCGTGATCG AAATGGCTAGAGAGAACCAGACCACTCAGAAGGGACAGAAGAACTCTAGGGAAAGGATGAAGAGGATCGAGGA AGGTATCAAAGAGCTTGGATCTCAGATCCTCAAAGAGCACCCTGTTGAGAACACTCAGCTCCAGAATGAGAAGCT CTACCTCTACTACCTCCAGAACGGAAGGGATATGTATGTGGATCAAGAGTTGGATATCAACAGGCTCTCTGATTAC GATGTTGATCATATCGTGCCACAGTCATTCTTGAAGGATGATTCTATCGATAACAAGGTGCTCACCAGGTCTGATA AGAACAGGGGTAAGAGTGATAACGTGCCAAGTGAAGAGGTTGTGAAGAAAATGAAGAACTATTGGAGGCAGCT CCTCAACGCTAAGCTCATCACTCAGAGAAAGTTCGATAACTTGACTAAGGCTGAGAGGGGAGGACTCTCTGAATT GGATAAGGCAGGATTCATCAAGAGGCAGCTTGTGGAAACCAGGCAGATCACTAAGCACGTTGCACAGATCCTCGA TTCTAGGATGAACACCAAGTACGATGAGAACGATAAGTTGATCAGGGAAGTGAAGGTTATCACCCTCAAGTCAAA GCTCGTGTCTGATTTCAGAAAGGATTTCCAATTCTACAAGGTGAGGGAAATCAACAACTACCACCACGCTCACGAT GCTTACCTTAACGCTGTTGTTGGAACCGCTCTCATCAAGAAGTATCCTAAGCTCGAGTCAGAGTTCGTGTACGGTG ATTACAAGGTGTACGATGTGAGGAAGATGATCGCTAAGTCTGAGCAAGAGATCGGAAAGGCTACCGCTAAGTATT TCTTCTACTCTAACATCATGAATTTCTTCAAGACCGAGATTACCCTCGCTAACGGTGAGATCAGAAAGAGGCCACTC ATCGAGACAAACGGTGAAACAGGTGAGATCGTGTGGGATAAGGGAAGGGATTTCGCTACCGTTAGAAAGGTGCT CTCTATGCCACAGGTGAACATCGTTAAGAAAACCGAGGTGCAGACCGGTGGATTCTCTAAAGAGTCTATCCTCCCT AAGAGGAACTCTGATAAGCTCATTGCTAGGAAGAAGGATTGGGACCCTAAGAAATACGGTGGTTTCGATTCTCCT ACCGTGGCTTACTCTGTTCTCGTTGTGGCTAAGGTTGAGAAGGGAAAGAGTAAGAAGCTCAAGTCTGTTAAGGAA CTTCTCGGAATCACTATCATGGAAAGGTCATCTTTCGAGAAGAACCCAATCGATTTCCTCGAGGCTAAGGGATACA AAGAGGTTAAGAAGGATCTCATCATCAAGCTCCCAAAGTACTCACTCTTCGAACTCGAGAACGGTAGAAAGAGGA TGCTCGCTTCTGCTGGTGAGCTTCAAAAGGGAAACGAGCTTGCTCTCCCATCTAAGTACGTTAACTTTCTTTACCTC GCTTCTCACTACGAGAAGTTGAAGGGATCTCCAGAAGATAACGAGCAGAAGCAACTTTTCGTTGAGCAGCACAAG CACTACTTGGATGAGATCATCGAGCAGATCTCTGAGTTCTCTAAAAGGGTGATCCTCGCTGATGCAAACCTCGATA AGGTGTTGTCTGCTTACAACAAGCACAGAGATAAGCCTATCAGGGAACAGGCAGAGAACATCATCCATCTCTTCAC CCTTACCAACCTCGGTGCTCCTGCTGCTTTCAAGTACTTCGATACAACCATCGATAGGAAGAGATACACCTCTACCA AAGAAGTGCTCGATGCTACCCTCATCCATCAGTCTATCACTGGACTCTACGAGACTAGGATCGATCTCTCACAGCTC GGTGGTGATTCAAGGGCTGATCCTAAGAAGAAGAGGAAGGTTggttcgTACCCATACGATGTTCCTGACTATGCGG GCTATCCCTATGACGTCCCGGACTATGCAGGATTGTATCCATATGACGTTCCAGATTACgccactagagctgcttaaGCTT CTCTAGCTAGAGTCGATCGACAAGCTCGAGTTTCTCCATAATAATGTGTGAGTAGTTCCCAGATAAGGGAATTAGG GTTCCTATAGGGTTTCGCTCATGTGTTGAGCATATAAGAAACCCTTAGTATGTATTTGTATTTGTAAAATACTTCTAT CAATAAAATTTCTAATTCCTAAAACCAAAATCCAGTACTAAAATCCAGAT Red font: CaMV 35S promoter with UBQ10, intron 1 (p35S); green font: bpNLS; purple font: SV40 NLS; black font: plant codon-optimized SpCas9 (pcoCas9) containing; dark blue font: Arabidopsis thaliana phosphoribosylanthranilate transferase (PAT1, Genbank no. M96073.1) intron 1 (Trp1 intron 1); orange font: 3xHA tag; blue font: CaMV 35S terminator (t35S).
❖ **Plant selection marker**: pNOS-NptII-tOCS cloned from pICSL11024 (pICH47732::NOSp-NPTII-OCST) (Addgene Plasmid #51144).

## References

Anzalone AV, Koblan LW, Liu DR (2020) Genome editing with CRISPR-Cas nucleases, base editors, transposases and prime editors. Nat Biotechnol 38 (7):824–844. doi:10.1038/s41587-020-0561-9

Ata H, Ekstrom TL, Martinez-Galvez G, Mann CM, Dvornikov AV, Schaefbauer KJ, Ma AC, Dobbs D, Clark KJ, Ekker SC (2018) Robust activation of microhomology-mediated end joining for precision gene editing applications. PLoS Genet 14 (9):e1007652. doi:10.1371/journal.pgen.1007652

Bae S, Kweon J, Kim HS, Kim JS (2014) Microhomology-based choice of Cas9 nuclease target sites. Nat Methods 11 (7):705–706. doi:10.1038/nmeth.3015

Brooks C, Nekrasov V, Lippman ZB, Van Eck J (2014) Efficient gene editing in tomato in the first generation using the clustered regularly interspaced short palindromic repeats/CRISPR-associated9 system. Plant Physiol 166 (3):1292–1297. doi:10.1104/pp.114.247577

Cermak T, Baltes NJ, Cegan R, Zhang Y, Voytas DF (2015) High-frequency, precise modification of the tomato genome. Genome Biol 16:232. doi:10.1186/s13059-015-0796-9

Dutta A, Eckelmann B, Adhikari S, Ahmed KM, Sengupta S, Pandey A, Hegde PM, Tsai MS, Tainer JA, Weinfeld M, Hegde ML, Mitra S (2017) Microhomology-mediated end joining is activated in irradiated human cells due to phosphorylation-dependent formation of the XRCC1 repair complex. Nucleic Acids Res 45 (5):2585–2599. doi:10.1093/nar/gkw1262

Hayashi A, Tanaka K (2019) Short-Homology-Mediated CRISPR/Cas9-Based Method for Genome Editing in Fission Yeast. G3 (Bethesda) 9 (4):1153-1163. doi:10.1534/g3.118.200976

Jinek M, Chylinski K, Fonfara I, Hauer M, Doudna JA, Charpentier E (2012) A programmable dual-RNA-guided DNA endonuclease in adaptive bacterial immunity. Science 337 (6096):816–821. doi:10.1126/science.1225829

Li P, Zhang L, Li Z, Xu C, Du X, Wu S (2019) Cas12a mediates efficient and precise endogenous gene tagging via MITI: microhomology-dependent targeted integrations. Cell Mol Life Sci. doi:10.1007/s00018-019-03396-8

Li XM, Chao DY, Wu Y, Huang X, Chen K, Cui LG, Su L, Ye WW, Chen H, Chen HC, Dong NQ, Guo T, Shi M, Feng Q, Zhang P, Han B, Shan JX, Gao JP, Lin HX (2015) Natural alleles of a proteasome alpha2 subunit gene contribute to thermotolerance and adaptation of African rice. Nat Genet 47 (7):827–833. doi:10.1038/ng.3305

Lin Q, Jin S, Zong Y, Yu H, Zhu Z, Liu G, Kou L, Wang Y, Qiu J-L, Li J, Gao C (2021) High-efficiency prime editing with optimized, paired pegRNAs in plants. Nature Biotechnology 39 (8):923–927. doi:10.1038/s41587-021-00868-w

Lu Y, Tian Y, Shen R, Yao Q, Wang M, Chen M, Dong J, Zhang T, Li F, Lei M, Zhu JK (2020a) Targeted, efficient sequence insertion and replacement in rice. Nat Biotechnol 38 (12):1402–1407. doi:10.1038/s41587-020-0581-5

Lu Y, Tian Y, Shen R, Yao Q, Zhong D, Zhang X, Zhu JK (2020b) Precise genome modification in tomato using an improved prime editing system. Plant Biotechnol J. doi:10.1111/pbi.13497

Matsuzaki S, Sakuma T, Yamamoto T (2024) REMOVER-PITCh: microhomology-assisted long-range gene replacement with highly multiplexed CRISPR-Cas9. In Vitro Cellular & Developmental Biology - Animal 60 (7):697–707. doi:10.1007/s11626-024-00850-1

Mazur BJ, Chui CF, Smith JK (1987) Isolation and characterization of plant genes coding for acetolactate synthase, the target enzyme for two classes of herbicides. Plant Physiol 85 (4):1110–1117. doi:10.1104/pp.85.4.1110

Nakade S, Tsubota T, Sakane Y, Kume S, Sakamoto N, Obara M, Daimon T, Sezutsu H, Yamamoto T, Sakuma T, Suzuki KT (2014) Microhomology-mediated end-joining-dependent integration of donor DNA in cells and animals using TALENs and CRISPR/Cas9. Nat Commun 5:5560. doi:10.1038/ncomms6560

Park J, Lim K, Kim J-S, Bae S (2016) Cas-analyzer: an online tool for assessing genome editing results using NGS data. Bioinformatics 33 (2):286–288. doi:10.1093/bioinformatics/btw561%JBioinformatics

Puchta H (2005) The repair of double-strand breaks in plants: mechanisms and consequences for genome evolution. J Exp Bot 56 (409):1–14. doi:10.1093/jxb/eri025

Robert F, Barbeau M, Ethier S, Dostie J, Pelletier J (2015) Pharmacological inhibition of DNA-PK stimulates Cas9-mediated genome editing. Genome Med 7:93. doi:10.1186/s13073-015-0215-6

Sakuma T, Nakade S, Sakane Y, Suzuki KT, Yamamoto T (2016) MMEJ-assisted gene knock-in using TALENs and CRISPR-Cas9 with the PITCh systems. Nat Protoc 11 (1):118–133. doi:10.1038/nprot.2015.140

Schindele A, Dorn A, Puchta H (2020) CRISPR/Cas brings plant biology and breeding into the fast lane. Curr Opin Biotechnol 61:7–14. doi:10.1016/j.copbio.2019.08.006

Sharma S, Javadekar SM, Pandey M, Srivastava M, Kumari R, Raghavan SC (2015) Homology and enzymatic requirements of microhomology-dependent alternative end joining. Cell Death Dis 6:e1697. doi:10.1038/cddis.2015.58

Tan J, Zhao Y, Wang B, Hao Y, Wang Y, Li Y, Luo W, Zong W, Li G, Chen S, Ma K, Xie X, Chen L, Liu YG, Zhu Q (2020) Efficient CRISPR/Cas9-based plant genomic fragment deletions by microhomology-mediated end joining. Plant Biotechnol J. doi:10.1111/pbi.13390

Villarreal DD, Lee K, Deem A, Shim EY, Malkova A, Lee SE (2012) Microhomology directs diverse DNA break repair pathways and chromosomal translocations. PLoS Genet 8 (11):e1003026. doi:10.1371/journal.pgen.1003026

Vu TV, Doan DTH, Tran MT, Sung YW, Song YJ, Kim J-Y (2021a) Improvement of the LbCas12a-crRNA System for Efficient Gene Targeting in Tomato. 12. doi:10.3389/fpls.2021.722552

Vu TV, Nguyen NT, Kim J, Hong JC, Kim JY (2024a) Prime editing: Mechanism insight and recent applications in plants. Plant Biotechnol J 22 (1):19–36. doi:10.1111/pbi.14188

Vu TV, Nguyen NT, Kim J, Song YJ, Nguyen TH, Kim JY (2024b) Optimized dicot prime editing enables heritable desired edits in tomato and Arabidopsis. Nat Plants. doi:10.1038/s41477-024-01786-w

Vu TV, Sivankalyani V, Kim E-J, Doan DTH, Tran MT, Kim J, Sung YW, Park M, Kang YJ, Kim J-Y (2020) Highly efficient homology-directed repair using CRISPR/Cpf1-geminiviral replicon in tomato. Plant Biotechnology Journal 18 (10):2133–2143. 10.1111/pbi.13373

Vu TV, Thi Hai Doan D, Kim J, Sung YW, Thi Tran M, Song YJ, Das S, Kim J-Y (2021b) CRISPR/Cas-based precision genome editing via microhomology-mediated end joining. Plant Biotechnology Journal 19 (2):230–239. 10.1111/pbi.13490

Wang M, Lu Y, Botella JR, Mao Y, Hua K, Zhu JK (2017) Gene Targeting by Homology-Directed Repair in Rice Using a Geminivirus-Based CRISPR/Cas9 System. Mol Plant 10 (7):1007–1010. doi:10.1016/j.molp.2017.03.002

Wolter F, Puchta H (2019) In planta gene targeting can be enhanced by the use of CRISPR/Cas12a. The Plant Journal 100 (5):1083–1094. 10.1111/tpj.14488

Woo JW, Kim J, Kwon SI, Corvalan C, Cho SW, Kim H, Kim SG, Kim ST, Choe S, Kim JS (2015) DNA-free genome editing in plants with preassembled CRISPR-Cas9 ribonucleoproteins. Nat Biotechnol 33 (11):1162–1164. doi:10.1038/nbt.3389

Yao X, Wang X, Liu J, Hu X, Shi L, Shen X, Ying W, Sun X, Wang X, Huang P, Yang H (2017) CRISPR/Cas9 - Mediated Precise Targeted Integration In Vivo Using a Double Cut Donor with Short Homology Arms. EBioMedicine 20:19–26. doi:10.1016/j.ebiom.2017.05.015

Yazdani M, Sun Z, Yuan H, Zeng S, Thannhauser TW, Vrebalov J, Ma Q, Xu Y, Fei Z, Van Eck J, Tian S, Tadmor Y, Giovannoni JJ, Li L (2019) Ectopic expression of ORANGE promotes carotenoid accumulation and fruit development in tomato. Plant Biotechnol J 17 (1):33–49. doi:10.1111/pbi.12945

Yuan H, Owsiany K, Sheeja TE, Zhou X, Rodriguez C, Li Y, Welsch R, Chayut N, Yang Y, Thannhauser TW, Parthasarathy MV, Xu Q, Deng X, Fei Z, Schaffer A, Katzir N, Burger J, Tadmor Y, Li L (2015) A Single Amino Acid Substitution in an ORANGE Protein Promotes Carotenoid Overaccumulation in Arabidopsis. Plant Physiol 169 (1):421–431. doi:10.1104/pp.15.00971

Zhang Z, Zeng W, Zhang W, Li J, Kong D, Zhang L, Wang R, Peng F, Kong Z, Ke Y, Zhang H, Kim C, Zhang H, Botella JR, Zhu J-K, Miki D (2022) Insights into the molecular mechanisms of CRISPR/Cas9-mediated gene targeting at multiple loci in Arabidopsis. Plant Physiology:kiac431. doi:10.1093/plphys/kiac431

Zhao Y, Thomas HD, Batey MA, Cowell IG, Richardson CJ, Griffin RJ, Calvert AH, Newell DR, Smith GC, Curtin NJ (2006) Preclinical evaluation of a potent novel DNA-dependent protein kinase inhibitor NU7441. Cancer Res 66 (10):5354–5362. doi:10.1158/0008-5472.CAN-05-4275

Zong Y, Liu Y, Xue C, Li B, Li X, Wang Y, Li J, Liu G, Huang X, Cao X, Gao C (2022) An engineered prime editor with enhanced editing efficiency in plants. Nature Biotechnology 40 (9):1394–1402. doi:10.1038/s41587-022-01254-w

